# A genus-specific nsp12 region impacts polymerase assembly in Alpha- and Gammacoronaviruses

**DOI:** 10.1101/2024.07.23.604833

**Authors:** Peter J. Hoferle, Thomas K. Anderson, Robert N. Kirchdoerfer

**Author notes:** These authors contributed equally to this work (Peter J. Hoferle, Thomas K. Anderson).

## Abstract

Coronavirus relevancy for human health has surged over the past 20 years as they have a propensity for spillover into humans from animal reservoirs resulting in pandemics such as COVID-19. The diversity within the *Coronavirinae* subfamily and high infection frequency in animal species worldwide creates a looming threat that calls for research across all genera within the *Coronavirinae* subfamily. We sought to contribute to the limited structural knowledge within the *Gammacoronavirus* genera and determined the structure of the viral core replication-transcription complex (RTC) from Infectious Bronchitis Virus (IBV) using single-particle cryo-EM. Comparison between our IBV structure with published RTC structures from other *Coronavirinae* genera reveals structural differences across genera. Using *in vitro* biochemical assays, we characterized these differences and revealed their differing involvement in core RTC formation across different genera. Our findings highlight the value of cross-genera *Coronavirinae* studies, as they show genera specific features in coronavirus genome replication. A broader knowledge of coronavirus replication will better prepare us for future coronavirus spillovers.

## Introduction

Coronaviruses belong to the *Nidovirales* order of positive-sense RNA viruses. Within *Nidovirales*, this diverse subfamily of viruses is divided into four genera: *Alpha*-, *Beta*-, *Gamma*- and *Deltacoronavirus*.(1) In 1931, the *Gammacoronavirus* infectious bronchitis virus (IBV) was the first coronavirus to be discovered.(2) Subsequently, additional members of the subfamily have been characterized including the human seasonal betacoronaviruses HKU1 and OC43 and alphacoronaviruses NL63 and 229E. (1) Since 2002, three animal betacoronaviruses have crossed into humans and caused disease outbreaks: SARS-CoV in 2002, MERS-CoV in 2012 and SARS-CoV-2 in 2019.(3–5) The emergence of SARS-CoV-2, the causative agent of COVID-19, led to a global pandemic that has resulted in large losses of life and significant burdens on both healthcare and the economy. In 2018, a recombinant canine-feline *Alphacoronavirus*, CCoV-HuPn-2018, was isolated from human patients hospitalized with pneumonia.(6) Although CCoV-HuPn-2018 is currently incapable of efficiently infecting humans, it is poised as a preemergent human pathogen.(7,8) The *Gamma*- and *Deltacoronavirus* genera contain numerous avian coronaviruses, and while no avian-to-human spillovers from these genera have been reported, the identification of recent independent infection of Haitian children with porcine deltacoroanvirus along with the consistent threat posed by avian viruses from the close contact between human and avian populations highlights the need for better characterization and monitoring of these animal coronaviruses.(9)

The *Gammacoronavirus* genus is subdivided into three subgenera: *Igacovirus, Brangacovirus* and *Cegacovirus*.(10) *Igacovirus* is currently recognized to have three species including *galli, pulli* and *anatis* (formerly duck coronavirus 2714). Isolates of IBV fall into both *galli* and *pulli* species while *anatis* is typically found in wild birds.(11) Infection of chickens with IBV typically initiates in the respiratory tract and some strains can additionally infect the reproductive tract and kidneys. Infection of the reproductive tract can lead to a decrease in egg quality while infection of the kidneys may lead to nephritis and death.(12,13) Respiratory tract infection may also weaken the immune system permitting secondary bacterial pneumonia.(14) Having a high prevalence in most parts of the world, IBV has been an immense economic burden on the poultry industry. Despite extensive vaccination campaigns against IBV, the large genetic diversity of the virus arising from mutation and recombination creates difficulties in providing broad protection from IBV infection.(15)

Coronavirus genomes encode a large number of structural and non-structural proteins used to replicate viral genomes, assemble new virions, and interact with the infected host cell.(16) The 5’ two-thirds of the viral genome encodes the viral non-structural proteins (nsps) responsible for viral RNA replication and transcription. These nsps are the products of polyprotein cleavage and are encoded within two open reading frames: ORF1a and ORF1b, with ORF1b accessed by -1 programmed ribosomal frameshifting at the end of ORF1a to produce either the pp1a or pp1ab polyproteins. Across coronavirus genera, ORF1a/b have similar organizations and cleavage products to assemble the necessary machinery for viral RNA synthesis. Within the functionally conserved suite of nsps, nsp12 encodes the RNA-dependent RNA polymerase as well as containing a second active site for a nucleotidyltransferase.(18, 19) For viruses of *Alpha*- and *Betacoronavirus*, nsp12 requires the replication factors nsp7 and nsp8 for robust RNA synthesis activity.(20, 21) These three nsps form the core polymerase complex that can perform processive RNA synthesis *in vitro*.(1) The majority of the work to characterize coronavirus polymerases has focused on betacoronaviruses leaving members of other genera relatively understudied.(22) A prior study of IBV polymerase has indicated an interaction of nsp12 with nsp8 though without a demonstration of polymerase activity or molecular descriptors.(23)

Structural studies of coronavirus polymerase complexes have largely focused on complexes from SARS-CoV-2 with limited polymerase structures from SARS-CoV and the *Alphacoronavirus* porcine epidemic diarrhea virus (PEDv).(21, 22, 24, 25) These structures have revealed similar nsp12 active site architectures and requirements for nsp7 and nsp8 replication factors. Expanding beyond this dataset dominated by structures of *Betacoronavirus* polymerases affords the opportunity to examine unique features of coronavirus polymerases across the subfamily while also identifying conserved mechanisms between these divergent viruses. Here, we use cryo-electron microscopy to solve the structure of the IBV polymerase complex, the first such polymerase structure from the *Gammacoronavirus* genus. We identified a genus-specific nsp12 loop that in PEDv and IBV contacts a subunit of nsp8. Subsequent biochemical analyses demonstrate the importance of this interaction and point to the potential of genus-specific polymerase assembly pathways. Continued investigation of coronavirus polymerases across this diverse group of viruses will aid in the development of broad-spectrum antiviral therapeutics and inform mechanisms for viral polymerase function.

## Results

### IBV shares replication factor requirements for RNA binding and synthesis with *Alpha*- and *Betacoronaviruses*

Recombinantly expressed and purified IBV nsp7, nsp8 and nsp12 combined with a fluorescently labeled RNA primer/template pair altered the mobility of the RNA on native-PAGE in a manner demonstrating that both nsp7 and nsp8 are required for RNA binding to nsp12 (**Fig. S4**). Adding nucleotides to this complex similarly demonstrated the IBV nsp12 requirement for both nsp7 and nsp8 for promoting robust RNA synthesis activity by primer extension assay (**Fig. 1A and S5**). The results of these assays indicate similar requirements for nsp7 and nsp8 for RNA binding and polymerase activity across the coronavirus subfamily.(20, 21, 25)

**Figure 1.**
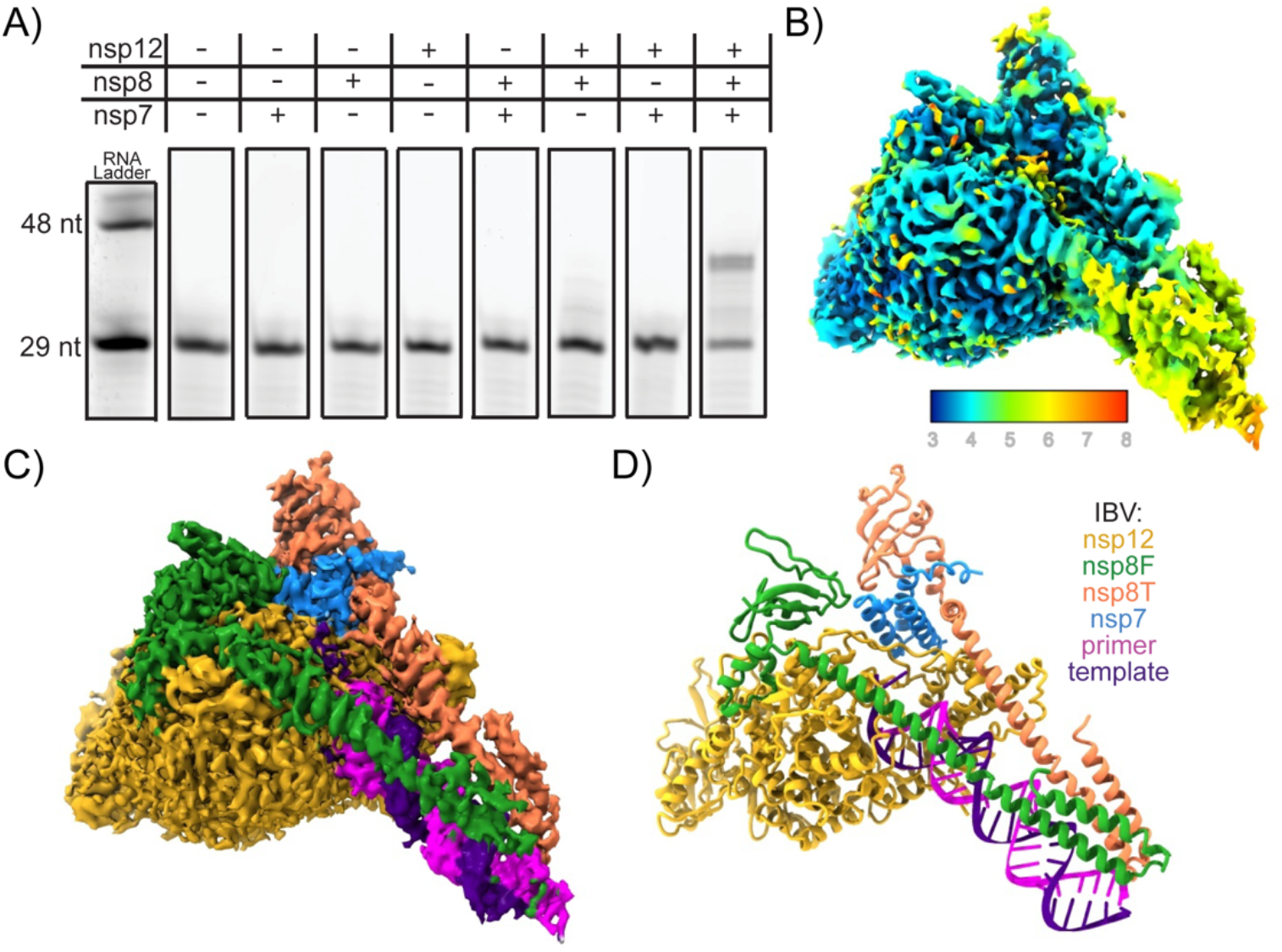
Structure of an active IBV polymerase complex: **A)** Extension of a short (29 nt) RNA primer to the length of the template RNA (38 nt) in the presence of the IBV polymerase complex. **B and C)** Cryo-EM reconstruction of the IBV polymerase complex colored by local resolution (**B**) or chain (**C**). **D)** Atomic model of the IBV polymerase complex built using the cryo-EM reconstruction.

### Structure of the IBV polymerase complex

Single-particle cryo-electron microscopy was used to solve the structure of the IBV nsp7-nsp8-nsp12-RNA complex. Our cryo-EM reconstruction has a resolution of 3.5 Å (**Fig. 1B,C, S2, S3 Table S1)**. Clearly visible in the map are densities for all components including nsp12, nsp7, two copies of nsp8, and the RNA substrate with a complex stoichiometry of 1:2:1 for nsp7:nsp8:nsp12 which is consistent with other coronavirus polymerase structures **(Fig. 1D)**.(21, 24, 25) Similar to *Alpha*- and *Betacoronavirus* polymerases one protomer of nsp8 binds the nsp12 fingers domain (nsp8_F_) while a second protomer binds to nsp12 as a nsp7-nsp8 heterodimer adjacent to the nsp12 thumb domain (nsp8_T_). The identification of nsp8_F_ is congruent with a previous biochemical study of IBV nsp12 demonstrating an interaction of nsp8 with nsp12 residues 1-400 which encompasses nearly the entirety of our observed nsp8_F_ binding site on nsp12.(23) As previously observed in coronavirus polymerase structures bound to duplex RNAs, the N-terminal extensions of each nsp8 form long helices to contact upstream double-stranded RNA extending from the polymerase active site.(21, 25) The IBV polymerase and nucleotidyltransferase active sites are well resolved and well conserved both in sequence and structure among SARS-CoV, SARS-CoV-2 and PEDv polymerases (SARS-CoV: 6NUR, SARS-CoV-2: 7KRP, PEDv: 8URB) suggesting the broad applicability of antiviral drugs targeting these sites.(26, 27)

### Structurally observed insertions and deletions in the *Gammacoronavirus* polymerase complex

Despite the high sequence and structural homology of coronavirus polymerases, we identified large insertions and deletions in nsp8 and nsp12 that result in unique conformations within the IBV polymerase complex structure. Many of these regions are distal to known active sites and protein-protein interfaces and their influence on the viral polymerase remains a topic for future study.

IBV nsp8 loop 173-181 contains an insertion not observed in other genera of coronaviruses **(Fig. S6)**. This loop sequence is well conserved among *Igacovirus* nsp8s while *Brangocovirus* nsp8s contain an additional three amino acid insertion and the *Cegacovirus* nsp8s have a nine amino acid deletion **(Fig. S7)**. *Betacoronavirus* nsp8s possess shorter loops in this nsp8 region with *Embecovirus* members, such as Murine Hepatitis Virus, having nsp8 loops nine amino acids shorter than IBV. Similarly, *Alpha*- and *Deltacoronavirus* nsp8s have loops five and 13 amino acids shorter than IBV, respectively. This IBV nsp8 region lacks secondary structure and forms an extended loop from the nsp8 C-terminal head domains, while in SARS-CoV, SARS-CoV-2 and PEDv this loop forms a pair of short helices **(Fig. S8)**. This loop is clearly visible in the IBV reconstructed density in both nsp8_F_ and nsp8_T_ though the density is weaker at the distal end of the loop particularly for nsp8_T_. Additional examination of IBV nsp8_T_ reveals an 18° rotation in the conformation of the nsp8 head domain relative to nsp7 when compared to corresponding domains from SARS-CoV-2 (24). This rotated nsp8_T_ head domain is similar to the orientation of nsp8_T_ for PEDv (20). IBV nsp8 122-129 is highly conserved among *Gammacoronavirus* though poorly conserved across *Coronavirinae* and in nsp8_T_ makes contacts with nsp7. IBV and PEDv nsp8_T_ regions 122-129 (IBV) make extensive contacts to nsp7 *α*2 while in the SARS-CoV and SARS-CoV-2 equivalent nsp8_T_ regions, the interactions are more restricted to nsp7 *α*3 **(Fig S6, S7, S9)**.(21, 24, 25)

In addition to nsp8, there are several insertions and deletions in IBV nsp12 particularly in the N-terminal nucleotidyltransferase domain.(19) There is a large insertion in IBV nsp12 loop 67-72 when compared to *Alpha*- and *Betacoronavirus* nsp12s (**Fig. S10)**. This loop sequence is well conserved in *Gammacoronavirus galli* and *pulli* with some length polymorphisms in other *Gammacoronavirus* nsp12s **(Fig. S11)**. In *Alpha*- and *Betacoronavirus* nsp12, this loop is four and eight amino acids shorter, respectively, and one to two amino acids longer in *Deltacoronavirus*. This loop is positioned on the opposite side of the nucleotidyltransferase domain from the enzyme active site and is distal to known protein-binding sites for nsp7, nsp8, nsp9, and nsp13 **(Fig. S12)**.(24, 28, 29)

There is a shortened loop in IBV nsp12 113-115 that is conserved in all *Gammacoronavirus* nsp12s except for *Cegacovirus* nsp12s which are one amino acid shorter, similar to *Deltacoronavirus* nsp12s **(Fig. S10, S11)**. In contrast, *Alpha*- and *Betacoronavirus* nsp12s are four amino acids longer in this region **(Fig. S10)**. This nsp12 loop lies within the nucleotidyltransferase domain but again is distant from the enzyme active site and known protein-binding sites **(Fig. S13)**.

IBV nsp12 contains a four amino acid insertion in loop 156-169 compared to *Alpha*-, *Beta*- and *Deltacoronavirus* nsp12s **(Fig. S10)**. In *Gammacoronavirus*, the nsp12 loop 156-169 (IBV) is conserved across *Gammacoronavirus galli* and *pulli* species while *Gammacoronavirus anatis* and *Brangacovirus* contain an additional 10 amino acid insertion with more divergent sequences **(Fig S11)**. *Cegacovirus* nsp12s have shorter loops of similar length to the other coronavirus genera (**Fig S10)**. Structurally, this region of nsp12 appears to be conserved in both *Alpha*- and *Betacoronavirus* nsp12s which is unsurprising given nsp12s’ moderate sequence conservation between these two genera. The insertion in IBV nsp12 loop 156-169 results in a shortened helical region and more extended loop region that extends outwards from the polymerase **(Fig. S14)**. While loop sequence insertions at this position have so far only been noted for coronaviruses infecting avian species, the lack of insertions here among avian viruses of *Deltacoronavirus* and the poor representation of *Cegacovirus* sequences that infect mammals in databases warns against identifying this loop as a potential host species determinant.

### Sequence and structural comparison of the IBV nsp12 loop 264-278

In the IBV nsp12 interface domain, amino acids 264-278 form a large loop that is in a dramatically different conformation when compared to the homologous nsp12 region in both *Alpha*- and *Betacoronavirus* **(Fig. 2B)**.(21, 25) This loop extends from the interface domain to contact the head domain of nsp8_F_. We define this nsp12 loop as being flanked by well conserved residues L279 and L280 that form the hydrophobic pocket for nsp8_F_ region 103-129 and nsp12 E263 that forms a well conserved salt bridge with K294 (IBV nsp12 numbering). The corresponding region in PEDv nsp12 (249-264) also extends to contact the nsp8_F_ head domain but adopts a different conformation to use distinct regions on both nsp12 and nsp8_F_ to form the protein-protein interaction.(21) In contrast, the corresponding loop in *Betacoronavirus* SARS-CoV and SARS-CoV-2 nsp12s extends away from the core complex not forming interactions with known replication factors.(24, 25)

**Figure 2.**
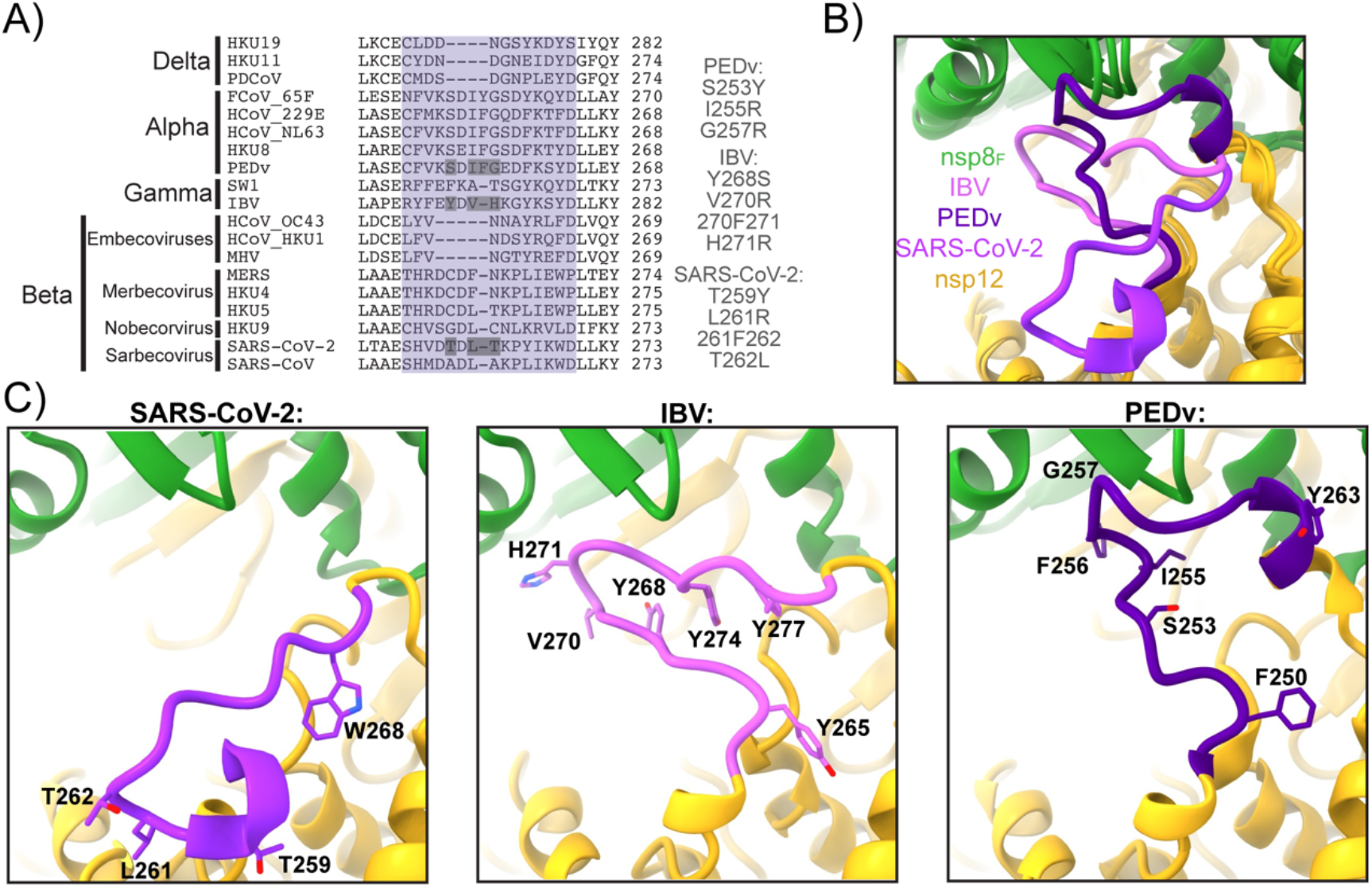
Altered sequence and structure of nsp12-nsp8_F_ interactions: **A)** Multiple sequence alignment of nsp12 residues 260-282 (IBV) across the coronavirus subfamily. The loop region of interest is highlighted in purple, and residues that were mutated in this study are listed to the right and have gray boxes around them. **B)** Superimposition of IBV, PEDv (8URB), and SARS-CoV-2 (6YYT) highlighting the altered conformations of the nsp12 loop (IBV residues 264-278). **C)** Individual snapshots of the nsp12 loop (IBV residues 264-278) from IBV (left), PEDv (center), and SARS-CoV-2 (right).

Examining sequence alignments for this region of nsp12, the *Betacoronavirus* subgenera *Nobecovirus, Sarbecovirus* and *Merbecovirus* have loop lengths similar to *Gammacoronavirus* nsp12 while *Embecovirus* nsp12s have loops that are four amino acids shorter (**Fig. 2A)**. While there is no structural data to provide insight into this nsp12 region for *Deltacoronavirus* polymerases, sequence alignment indicates a shortening of this loop by three amino acids relative to IBV. Hence, we hypothesize that the nsp12 loop does not contact nsp8_F_ in *Deltacoronavirus* nsp12s. In contrast *Alphacoronavirus* nsp12s have a one amino insertion, adding a Phe at position 256 (PEDv numbering). PEDv nsp12 F256 lies at the apex of the nsp12 loop and packs into the hydrophobic surface between nsp12 and nsp8_F_. To test the role of F256 in PEDv complex assembly we produced a recombinant PEDv nsp12 with F256 deleted (PEDv nsp12 ΔF256) which resulted in nearly a complete loss of polymerase activity using our aforementioned *in vitro* RNA primer extension assay (**Fig. 3)**. Complementary to this PEDv nsp12 deletion, we inserted a Phe into the corresponding nsp12 positions of IBV (270F271) and SARS-CoV-2 (261F262). Neither of these insertions diminished the ability of the mutant polymerases to bind RNA or extend primers (**Fig. 3 and S15, S16)**. In IBV nsp12, while this loop contacts nsp8_F_, the 270F271 insertion would be expected to be surface exposed owing to different utilization of this nsp12 region to contact the nsp8_F_ head domain.

**Figure 3.**
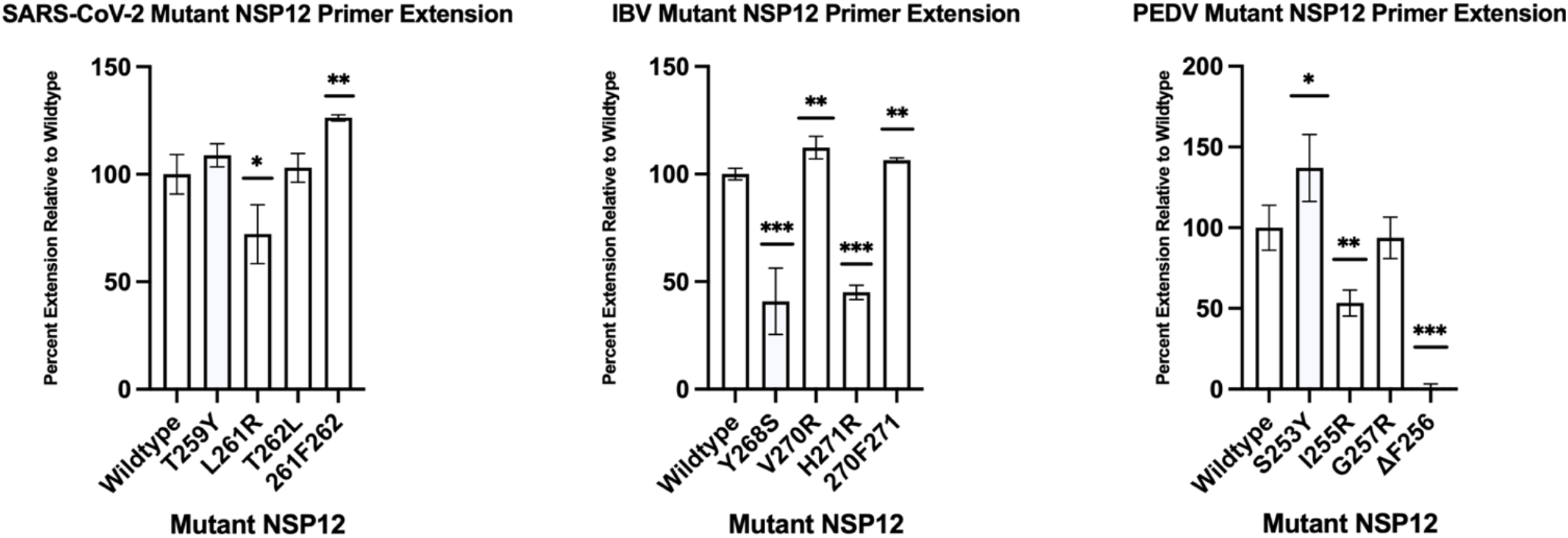
Mutant nsp12 primer extensions: Point mutations and insertions of nsp12 from SARS-CoV-2 (left), IBV (center), and PEDv (right) were tested for their ability to form the core-RTC and extend RNA *in vitro*. For each coronavirus, results are presented as % activity of the polymerase complex with respect to wildtype nsp12. Error bars indicate standard deviation of the triplicates. Using an unpaired t-test comparing each mutant to wildtype, ^“*”^ denotes *P* < 0.05, ^“**”^ denotes *P* < 0.001, and ^“***”^ denotes *P* < 0.0001.

Examining the conformation of the IBV nsp12 264-278 loop, Tyr residues at 268, 274 and 277 undergo aromatic stacking and likely stabilize this unique loop conformation. To test the importance of the tyrosine stacking in maintaining the IBV nsp12 264-278 loop conformation, we created an IBV nsp12 with a Y268S mutation. IBV nsp12 Y268S showed a more than 50% reduction in polymerase activity as well as a strong defect in RNA binding (**Fig. 3 and S15**,**16)**. These defects in polymerase activities likely are caused by a failure to properly assemble IBV nsp8_F_ on nsp12 Y268S and point to the importance of this IBV nsp12 loop region for polymerase complex binding to RNA. Complementary mutations in PEDv nsp12 (S253Y) or SARS-CoV-2 nsp12 (T259Y) did not have major effects on viral polymerase activities supporting the observed distinct conformations and interactions of these nsp12 loop regions (**Fig. 3 and Fig S15**,**16)**.

The presence of an aromatic residue at nsp12 277 (IBV numbering, SARS-CoV-2 W268, PEDv Y263) is well conserved across the coronavirus subfamily, however, these aromatic residues are placed in very different contexts within this loop region. In PEDv, nsp12-Y263 is surface exposed while in IBV nsp12-Y277 participates in the tyrosine stacking interaction that stabilizes this unique loop conformation. In contrast, SARS-CoV and SARS-CoV-2 nsp12-W268 is oriented into a hydrophobic pocket on nsp12 which may drive the diversion of this loop into the observed outwards directed conformation. Structural comparison of the SARS-CoV-2 nsp12-W268 hydrophobic pocket shows that the homologous pocket is occupied by IBV nsp12-Y265 or PEDv-F250 **(Fig. 2)**. The functional constraint of needing to insert an aromatic residue into this hydrophobic pocket may be driving loop conformational differences and presentation of this nsp12 loop to nsp8_F_ in IBV and PEDv polymerase complexes.

To further examine specific interactions at the IBV nsp12 264-278 and PEDv nsp12 249-264 loop apexes with their respective nsp8_F_ head domains, we generated nsp12 mutants for key residues in the protein interfaces and complementary mutations in other coronavirus polymerases (**Fig. 3 and S15**,**16)**. In addition to PEDv nsp12 F256, I255 contributes to the buried hydrophobic surface between nsp12 and the nsp8_F_ head domain. A PEDv nsp12-I255R mutation reduced primer extension activity by 50% and prevented strong RNA binding to the PEDv polymerase complex. Homologous mutations to IBV nsp12 (V271R) and SARS-CoV-2 nsp12 (L261R) did not have large effects on either polymerase complex. For IBV nsp12, H271 resides at the apex of the 264-278 loop to contact the nsp8_F_ head domain. IBV nsp12-H271R had a 50% reduction in polymerase activity and a significant loss in RNA binding activity. Mutations to homologous positions in PEDv nsp12 (G257R) or SARS-CoV-2 nsp12 (T262L) had no effect on either polymerase primer extension or RNA binding activity. These targeted mutations highlight the distinct interactions of each genera’s nsp12 loop with nsp8_F_ and that disrupting these interactions has genus-specific negative impacts on polymerase activity.

## Discussion

Here we have presented the first structure of a *Gammacoronavirus* polymerase complex showing the IBV RNA polymerase bound to its essential replication factors and RNA. Structural comparisons highlight a loop in IBV nsp12 (residues 264-278) that is in an alternate conformation than previous polymerase complex structures from PEDv and SARS-CoV-2. Mutagenesis of key residues in this protein region among these three polymerase complexes were a detriment to IBV and PEDv polymerase RNA-binding and primer extension activities. The inability of these polymerases to bind RNA is likely a result of defects in the nsp12s’ ability to assemble properly with replication factor nsp8_F_. It has been previously shown that replication factors nsp7 and nsp8 are essential for nsp12 RNA binding and that disruptions to the nsp8_F_ head domain - nsp12 interaction resulted in polymerases incapable of extending primers or binding RNA despite not fully blocking nsp8_F_ subunit binding to the complex.(20, 21) We identify this nsp12 loop as a genus-specific structural feature that is functionally important for the proper assembly of *Alpha*- and *Gammacoronavirus* polymerases. Work on SARS-CoV-2 polymerase complexes has identified the binding of nsp8_F_ as a rate limiting step in polymerase assembly.(30) The observed altered interactions of IBV and PEDv nsp12 with nsp8_F_ may indicate the existence of alternate polymerase assembly pathways across diverse viruses. Our observed structural differences among viral nsps with functional consequences for polymerase activity highlight the need to consider alternate assembly and functional pathways across diverse coronavirus genera more broadly. Interestingly, the IBV nsp12 264-278 and PEDv nsp12 249-264 loops neighbor SARS-CoV-2 nsp12 P323. During the early months of the COVID-19 pandemic variant strains carrying a nsp12 P323L mutation along with a D614G mutation in the viral spike rapidly rose to prominence.(31) The impacts of the SARS-CoV-2 nsp12 P323L mutation on polymerase activity remain unclear but the spatial proximity of this mutation to the observed altered loop conformations in IBV and PEDv polymerases creates the possibility that this region of the nsp12 polymerase has a role in modulating the activity and assembly of the coronavirus polymerase complex across viral evolution.

## Materials and Methods

### DNA Constructs

All IBV, PEDv, and SARS-CoV-2 nsp gene sequences were codon optimized (Genscript). Sequences for *Gammacoronavirus galli* IBV/M41/Y28 proteins originate from the GenBank sequence QWC71293.1. SARS-CoV-2 protein sequences originate from the GenBank sequence UHD90671.1. PEDv protein sequences originate from the GenBank sequence AKJ21892.1. IBV nsp7 and PEDv nsp7 were cloned into the pET46 vector with C-terminal TEV protease cleavage site and hexahistidine tag. IBV nsp8 was cloned into pET45 vector with an N-terminal hexahistidine tag and TEV protease cleavage site. IBV nsp12, PEDv nsp8 and SARS-CoV-2 nsp7 and nsp8 were cloned into pET46 vectors with N-terminal hexahistidine tags and TEV protease cleavage sites. SARS-CoV-2 and PEDv nsp12 were cloned into pFastBac vectors with C-terminal TEV cleavage site and Strep II tags. Mutant nsp12 vectors were made by performing site-directed mutagenesis on the wildtype nsp12 vectors. The sequences of all open reading frames in plasmids were confirmed using Sanger sequencing.

### Protein Expression

Nsp7 and nsp8 were expressed in Rosetta 2pLysS *Escherichia coli* (*E. coli*) cells (Novagen). Cultures were grown at 37°C until they reached an OD_600_ of 0.6-0.8 where they were induced with IPTG (isopropyl β-D-1-thiogalactopyranoside) at a final concentration of 500 µM and incubated overnight at 16°C. Bacterial cells were pelleted and resuspended in wash buffer (10 mM Tris-Cl, 300 mM sodium chloride, 30 mM imidazole, 2 mM dithiothreitol (DTT), pH 8). Cells were then lysed using a microfluidizer (Microfluidics) and lysate was cleared using centrifugation and filtration. Lysate supernatant was used to batch bind to Ni-NTA beads (Qiagen) for 30 minutes before loading onto a gravity column. Beads were washed with wash buffer, then protein was eluted from beads using elution buffer (10 mM Tris-Cl, 300 mM sodium chloride, 300 mM imidazole, 2 mM DTT, pH 8). Eluted proteins were buffer exchanged by dialysis (10 mM TrisCl, 300 mM sodium chloride, 2 mM DTT, pH 8) while cleaving off the tag with tobacco etch viral (TEV) protease (1% w/w) at 4°C overnight. Proteins were passed back over a Ni-NTA column, collecting the flowthrough containing the cleaved protein sample. Protein was concentrated, then loaded onto a Superdex 200 10/300 Increase GL column (Cytiva) for size exclusion (25 mM Tris-Cl, 300 mM sodium chloride, 2 mM DTT, pH 8). Protein peak fractions were pooled and concentrated, then aliquoted and flash-frozen with liquid nitrogen. Proteins were stored at - 80°C until use. See also **Fig. S1**.

IBV nsp12 was expressed and purified using the same protocol as above but with alternate buffers. Ni-NTA wash buffer contained 25 mM sodium-HEPES, 300 mM sodium chloride, 30 mM imidazole, 1 mM magnesium chloride, 2 mM DTT, pH 7.5. Ni-NTA elution buffer contained 10 mM sodium-HEPES, 300 mM sodium chloride, 300 mM imidazole, 1 mM magnesium chloride, 2 mM DTT, pH 7.5. Dialysis buffer contained 10 mM sodium-HEPES, 300 mM sodium chloride, 1 mM magnesium chloride, 2 mM DTT, pH 7.5. Size exclusion buffer contained 25 mM sodium-HEPES, 300 mM sodium chloride, 100 µM magnesium chloride, 2 mM tris(2-carboxyethyl)phosphine (TCEP), pH 7.5. See also **Fig. S1**.

PEDv and SARS-CoV-2 nsp12 pFastBac vectors were transformed into DH10Bac *E. coli* to generate recombinant Bacmid plasmids. Bacmid plasmids were transfected into Sf9 cells to produce baculovirus stocks that were then amplified twice before being used to infect Sf21 cells. After two days of incubation at 27°C, infected Sf21 cells were pelleted, resuspended in wash buffer (25 mM sodium-HEPES, 300 mM sodium chloride, 1 mM magnesium chloride, 5 mM DTT, pH 7.4) with an added 143 uL of BioLock (IBA Lifesciences) and lysed with a microfluidizer. Lysed cells were cleared via centrifugation and filtration. Lysates were bound to streptactin beads (IBA Lifesciences) in batch for 30 minutes then loaded onto a gravity column. Beads were washed with wash buffer and proteins eluted with elution buffer (strep wash buffer with additional 2.5 mM desthiobiotin). Proteins were then concentrated, and further purified via SEC on a Superdex 200 10/300 Increase GL column with SEC buffer (25 mM HEPES, 300 mM sodium chloride, 100 µM magnesium chloride, 2 mM TCEP, pH 7.4). Protein peak fractions were pooled, concentrated, aliquoted and then flash-frozen with liquid nitrogen. Aliquoted samples were stored at -80°C until use. See also **Fig. S1**.

### RNA Substrate Preparation

RNA primers with 5’ fluorescein tags (6-FAM) were annealed to longer template RNA substrates. Formation of duplex RNAs was carried out in RNA annealing buffer (2.5 mM potassium chloride, 2.5 mM HEPES, 0.5 mM magnesium chloride, pH 7.4), with a primer:template ratio of 1:1.2. After mixing, samples were heated at 95°C for 5 minutes, then slowly cooled until reaching room temperature. Annealed substrates were used immediately or stored at -20°C.

RNA Primer for *in vitro* assays:

5’ – CAUUCUCCUAAGAAGCUAUUAAAAUCACA– 3’

RNA Template for *in vitro* assays:

5’ – AAAAAGGGUUGUGAUUUUAAUAGCUUCUUAGGAGAAUG– 3’

RNA Primer for structure determination:

5’ – CAUUCUCCUAAGAAGCUAUUAAAAUCACAGAUU– 3’

RNA Template for structure determination:

5’ – CAGUGUCAUGGAAAAACAGA*AA*AAUCUGUGAU UUUAAUAGCUUCUUAGGAGAAUG– 3’

### Primer Extension

Primer extension assays were carried out in 20 µL volumes with final buffer concentrations of 10 mM Tris-Cl pH 8, 2 mM magnesium chloride, 1 mM DTT and either 10 mM sodium chloride (PEDv and SARS-CoV-2) or 100 mM K-Glu (IBV). Nsp7 and nsp8 replication proteins and nsp12 were combined at final concentrations of 1.5 µM and 500 nM respectively. After combining, proteins were incubated together at room temperature for 15 minutes followed by the addition of duplex RNA substrate to 250 nM. After another 15-minute incubation at room temperature, 500 µM of each ribonucleotide was added to begin the extension. Reaction conditions were specific to each viral polymerase complex: 1 min at room temperature for SARS-CoV-2, 1 min at 30°C for PEDv, 30 min at 30°C for IBV. Reactions were then quenched by adding two volumes of denaturing RNA gel loading buffer (95% formamide (v/v), 2 mM EDTA, and 0.75 mM bromophenol blue). Quenched reactions were heated at 95°C for 15 minutes, then loaded on a denaturing urea-PAGE gel (8 M urea, 15% polyacrylamide) and run in TBE running buffer (89 mM TrisCl, 89 mM boric acid, 2 mM EDTA, pH 8.3). Gels were imaged using a GE Typhoon FLA 9000 scanner, using FAM tag excitation at 470 nm and measuring emission at 530 nm. The bands were analyzed using ImageJ.(32)

### Electrophoretic mobility shift assay

Electrophoretic mobility shift assays (EMSA) were carried out in 20 µL reaction volumes in buffer conditions of 10 mM Tris-Cl pH 7.4, 20 mM magnesium chloride, 1 mM DTT and either 10 mM sodium chloride (PEDv and SARS-CoV-2) or 10 mM potassium glutamate (IBV). Proteins were combined at final concentrations of nsp7 (3 µM), nsp8 (3 µM), and nsp12 (1 µM). Viral proteins were first mixed and allowed to incubate at room temperature for 15 minutes. RNA substrate was then added (250 nM final conc.), and the reaction incubated for an additional 15 minutes at room temperature. Finally, 10X non-denaturing gel loading buffer (10 mM Tris-Cl, 1 mM EDTA, 50% (v/v) glycerol, 0.75 mM bromophenol blue) was added to the reaction. Samples were run on a 4.5% non-denaturing PAGE gel with IBV and SARS-CoV-2 run on TBE native-PAGE gels while PEDV was run on clear native (CN)-PAGE gels.(33) Gels were scanned using a Typhoon imager scanning for FAM fluorescence. Bands were quantitated using ImageJ.(32)

### Specimen Preparation for cryoEM

IBV complexes were initially assembled at a total protein concentration of 2 mg/mL in cryoEM freezing buffer (10 mM Tris-Cl pH 8, 100 mM potassium glutamate, 2 mM MgCl_2_, and 1 mM DTT). Proteins and RNA were mixed at a ratio of 2:3:1:1.2 nsp7:nsp8:nsp12:RNA. To assemble the complexes, proteins were diluted in freezing buffer then combined and incubated at 25°C for 15 minutes before RNA was added and incubated for another 15 minutes at 25°C. After assembly, complexes were concentrated to 4 mg/mL total protein using ultrafiltration with a 100 kDa molecular weight cutoff. Samples were stored on ice prior to grid freezing.

Samples were frozen on UltraAuFoil R1.2/1.3 300 mesh grids (Quantifoil) using a Vitrobot Mark IV (ThermoFisher Scientific). Grids were freshly glow discharged using a GloQube Plus (Quorom) for 20 seconds with a current of 20 mA in an air atmosphere, creating a negative surface charge. Immediately prior to blotting, 0.5 µL of 42 mM 3-([3-cholamidopropyl] dimethylammonio)-2-hydroxy-1-propanesulfonate (CHAPSO) was added to 3 µL of sample. 3 µL of sample with CHAPSO (6 mM final) was spotted onto grids before double-sided blotting and vitrification in liquid ethane. Vitrobot chamber conditions were set to 100% humidity and 4°C.

### CryoEM Data Collection, Processing, and Model Building

EPU (ThermoFisher Scientific) was used for data collection on a Talos Arctica 200 keV transmission electron microscope (ThemoFisher Scientific). Movies were collected using a K3 direct electron detector (Gatan) in CDS mode. A GIF quantum energy filter was used with a slit width of 20 eV. Data was collected with no stage tilt at a magnification of 79,000x with a pixel size of 1.064 Å, and a defocus range of -0.5 to -2.0 μm with a step size of 0.5 µm. Total dose per movie was 60 e^-^/Å^2^.

Data were processed using cryoSPARC v4.3.0.(34) After patch motion correction and CTF estimation, 2,633,255 particles were picked using blob picker and extracted at a box size of 256 pixels. Particles were subjected to multiple rounds of 2D classification before three *ab initio* models were generated. Particles were classified by heterogeneous refinement using the three *ab initio* models as initial models. Output maps and classified particle stacks from heterogenous classification were used as inputs for non-uniform refinement. Particles from the class that resembled a polymerase complex were further classified by producing two *ab initio* models that were then used for heterogeneous refinement. The final reconstruction was produced using non-uniform refinement with 179,183 particles **(Fig S2**,**3, Table S1)**.

To build a starting coordinate model, we used AlphaFold 2 to create the IBV nsp12 RdRP and docked nsp7, nsp8, and duplex RNA from model 6YYT.pdb into our cryoEM reconstruction using ChimeraX.(25, 35–37) Model building and sequence changes were performed in Coot.(38) Iterative real-space refinement in Phenix and model building and adjustments using both ISOLDE and Coot were done to generate the final coordinate model.(38–40)

## Data Availability

The IBV core polymerase electron density map has been deposited in the Electron Microscopy Data Bank (EMDB: 45805) and the complex model has been deposited in the Protein Data Bank (PDB: 9CPO). All other data can be found within the manuscript and supporting information document.

## Supporting Information

This article contains supporting information.

## Funding and Additional Information

This work was supported by NIH/NIAID AI158463 to R.N.K.

