## Supplemental tables and figures for "A genus-specific nsp12 region impacts polymerase assembly in Alpha- and Gammacoronaviruses"

**Affiliations**
[1]Department of Biochemistry, Institute for Molecular Virology, Center for Quantitative Cell Imaging, University of Wisconsin-Madison, Madison, Wisconsin 53706

*These authors contributed equally to this work

Supplemental Table S1
Supplemental Figures S1-16

| | |
|---|---|
| EMDB | 45805 |
| PDB | 9CPO |
| Microscope | Talos Arctica |
| Voltage (kV) | 200 |
| Detector | K3 direct electron detector (Gatan) |
| Dose Rate ($e^-$/pixel/sec) | 14.2 |
| Exposure Time (sec) | 4.78 |
| Electron Exposure ($e^-/Å^2$) | 60 |
| Frames (no.) | 60 |
| Defocus Values (µm) | -0.5, -1.0, -1.5, -2.0 |
| Data Collection Mode | EFTEM, Counting, CDS |
| Nominal Magnification | 79,000 |
| Pixel Size (Å) | 1.064 |
| Symmetry Imposed | C1 |
| Movies Collected (no.) | 5,777 |
| Initial Particle Images (no.) | 2,633,225 |
| Final Particle Images (no.) | 179,183 |
| Map Resolution (Å) – GSFSC | 3.5 |
| Initial Models Used (PDB ID) | 6YYT |
| Non-hydrogen Atoms | 12,465 |
| Protein Residues | 1,455 |
| Nucleic Acid Residues | 63 |
| Other Atoms | 2 $Zn^{2+}$ |
| R.M.S. Deviations | |
| Bond Lengths (Å) | 0.003 |
| Bond angles (°) | 0.463 |
| MolProbity Score | 1.60 |
| Clashscore | 6.61 |
| Ramachandran Plot | |
| Favored (%) | 96 |
| Allowed (%) | 4 |
| Disallowed (%) | 0 |

**Table S1, cryo-EM data collection and refinement:** Information provided is for the cryoEM data collection, and processing that produced the electron density map for the IBV polymerase complex. PDB and EMDB codes are not provided as final model adjustments for PDB submission are still being done, in addition, because of this model validation statistics may be slightly altered compared after publication.

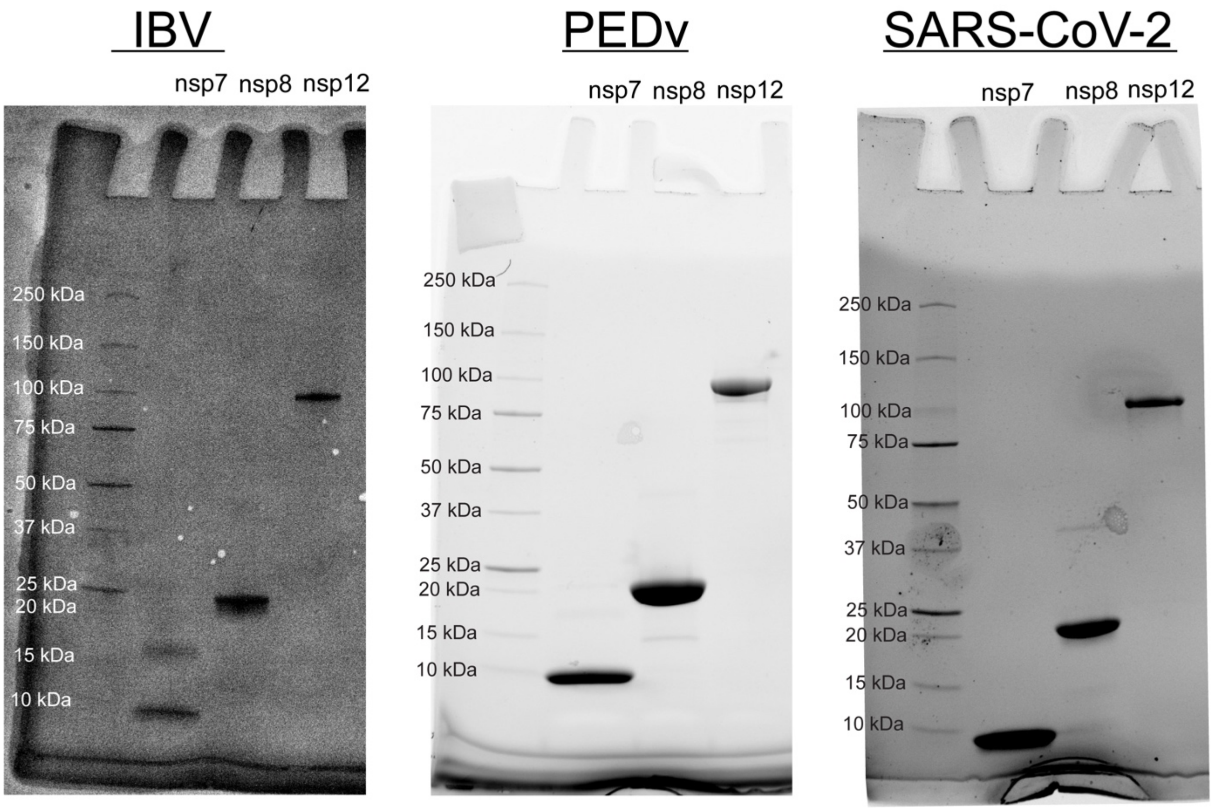


**Figure S1 SDS-PAGE analysis of viral RTC proteins:** Purified recombinant viral proteins used for activity assays and structure determination analyzed by SDS-PAGE. IBV proteins were visualized by Coomasssie staining while gels for PEDv and SARS-CoV-2 were visualized using UV fluorescence on stain-free gels (BioRad). Each gel was run with ladder on far-left lane with molecular weights labeled. Expected MW of each protein is as follows: nsp7 – ~9 kDa, nsp8 – ~22 kDa, nsp12- ~108 kDa.

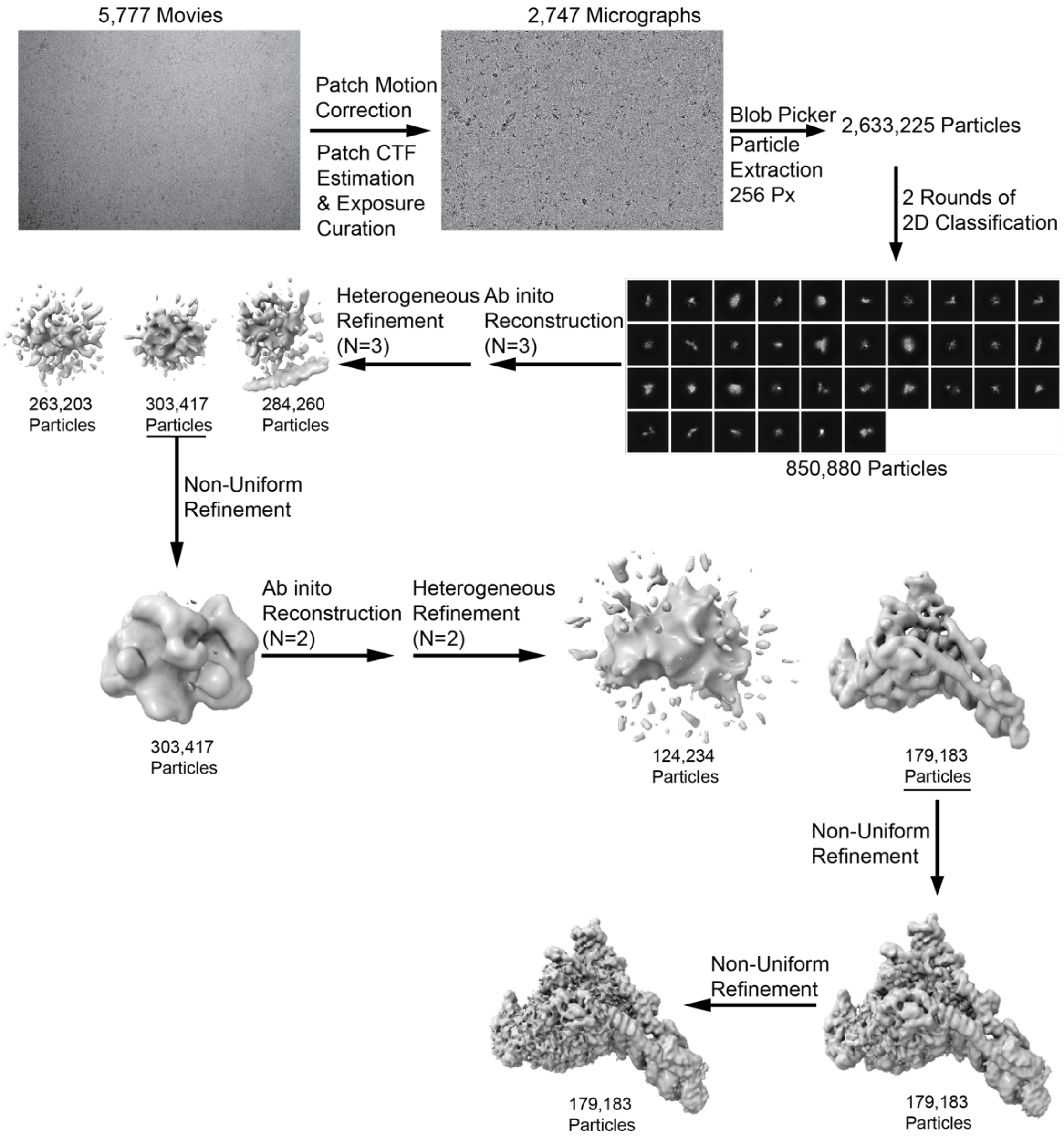


**Figure S2 Cryo-EM data processing pipeline:** Workflow for EM data processing using Cryo-Sparc. 5,777 movies were processed to micrographs and subsequently to individual particles which were filtered through two rounds of 2D classification. Ab initio reconstruction followed by heterogenous refinement was done twice to finalize on data set with 179,183 particles. Two rounds of non-uniform refinement were performed to give final refined 3.5 angstrom map.

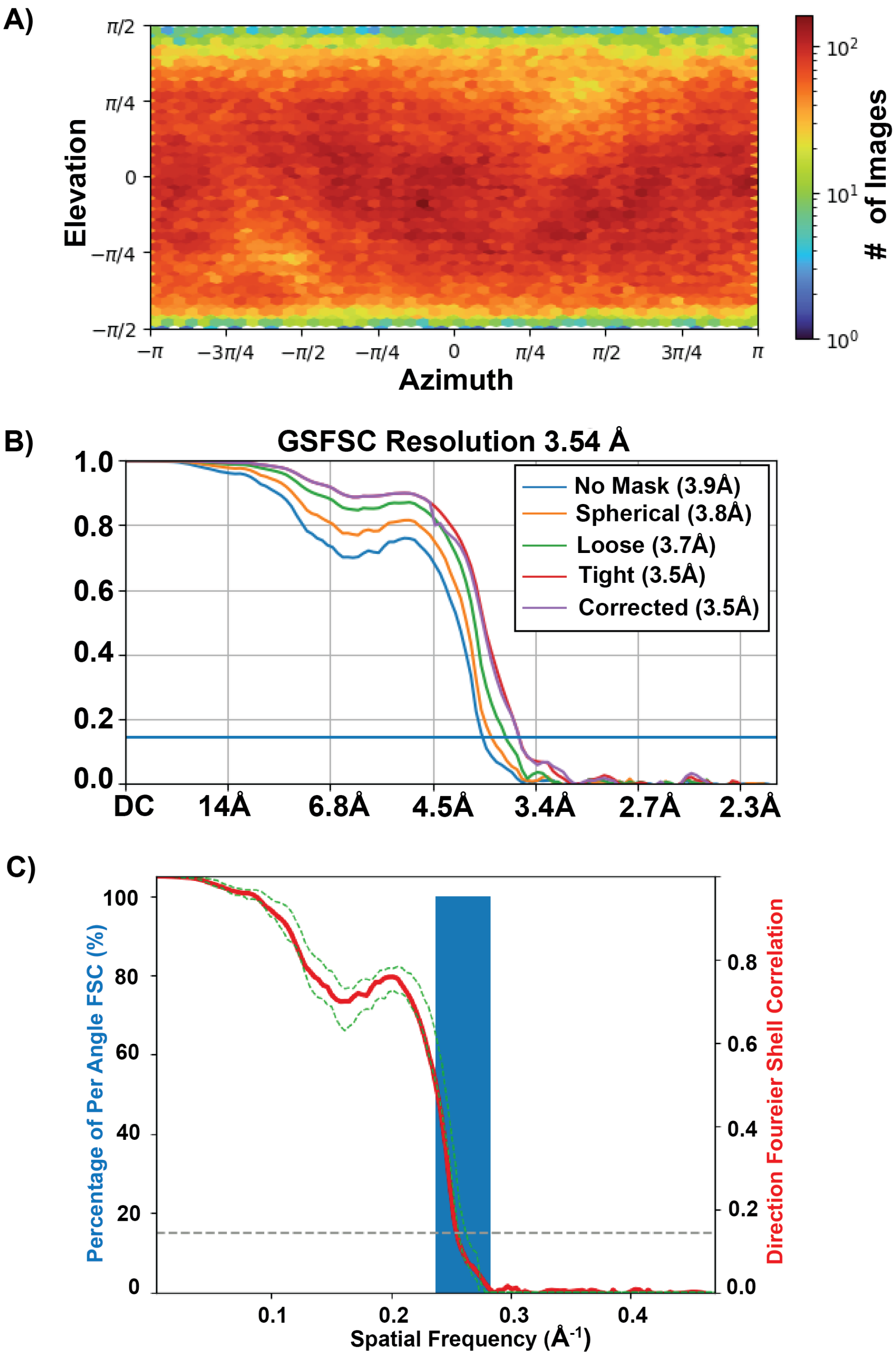


**Figure S3 Cryo-EM data validation:** A) Particle orientation distribution of for the final reconstructed EM map. B) Gold standard Fourier shell correlation plot from cryo-EM data. Blue line represents 0.143 cutoff. C) 3D Fourier shell correlation for the final refinement. Sphericity is 0.983 out of 1 and the unmasked global resolution is 3.95 angstrom.

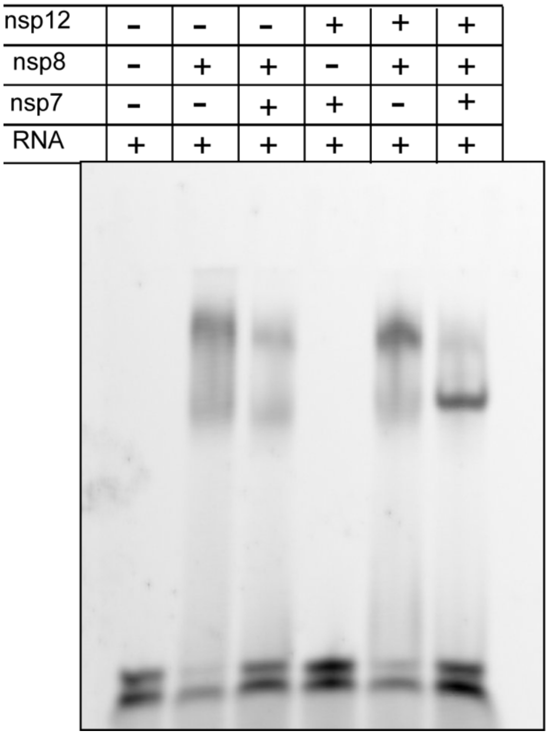


**Figure S4 Native-PAGE IBV RTC RNA substrate affinity:** Native PAGE gel analyzing IBV RTC binding to FAM tagged RNA duplex. Piecewise controls used to differentiate individual nsp vs whole complex interaction with RNA.

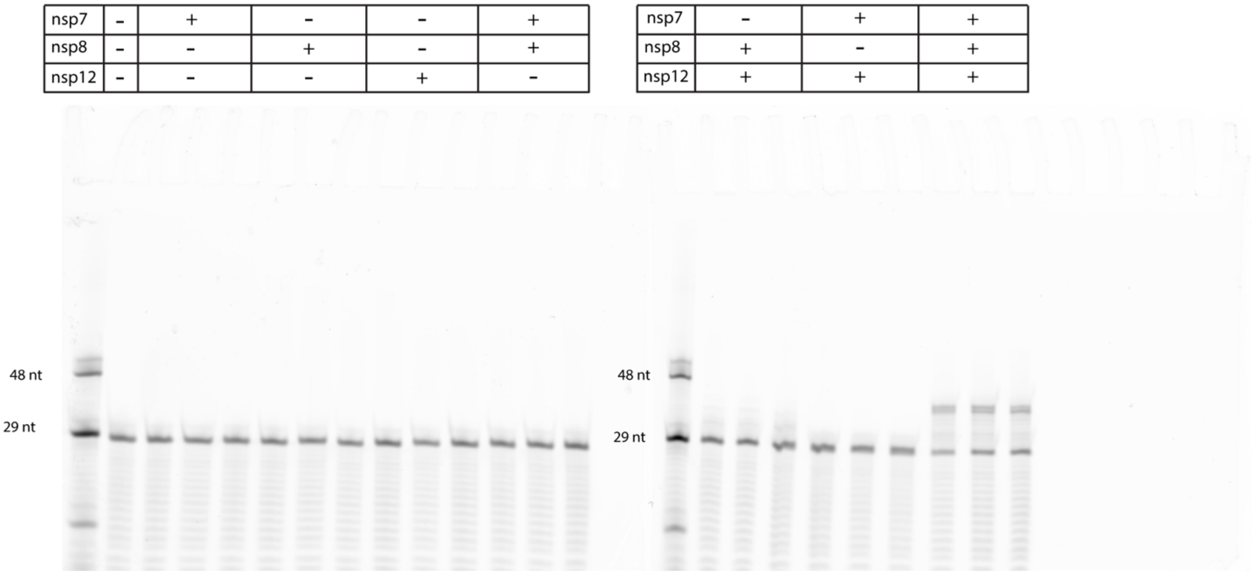


**Figure S5 IBV RTC Activity Assay:** Primer extension assay designed to assess activity of RTC based on ability to extend FAM tagged 29 nt RNA primer to the length of the 38 nt template. Triplicates ran with various combination of nsp7,8, and 12 mixed with RNA substrate. RNA analyzed on urea-PAGE gel.

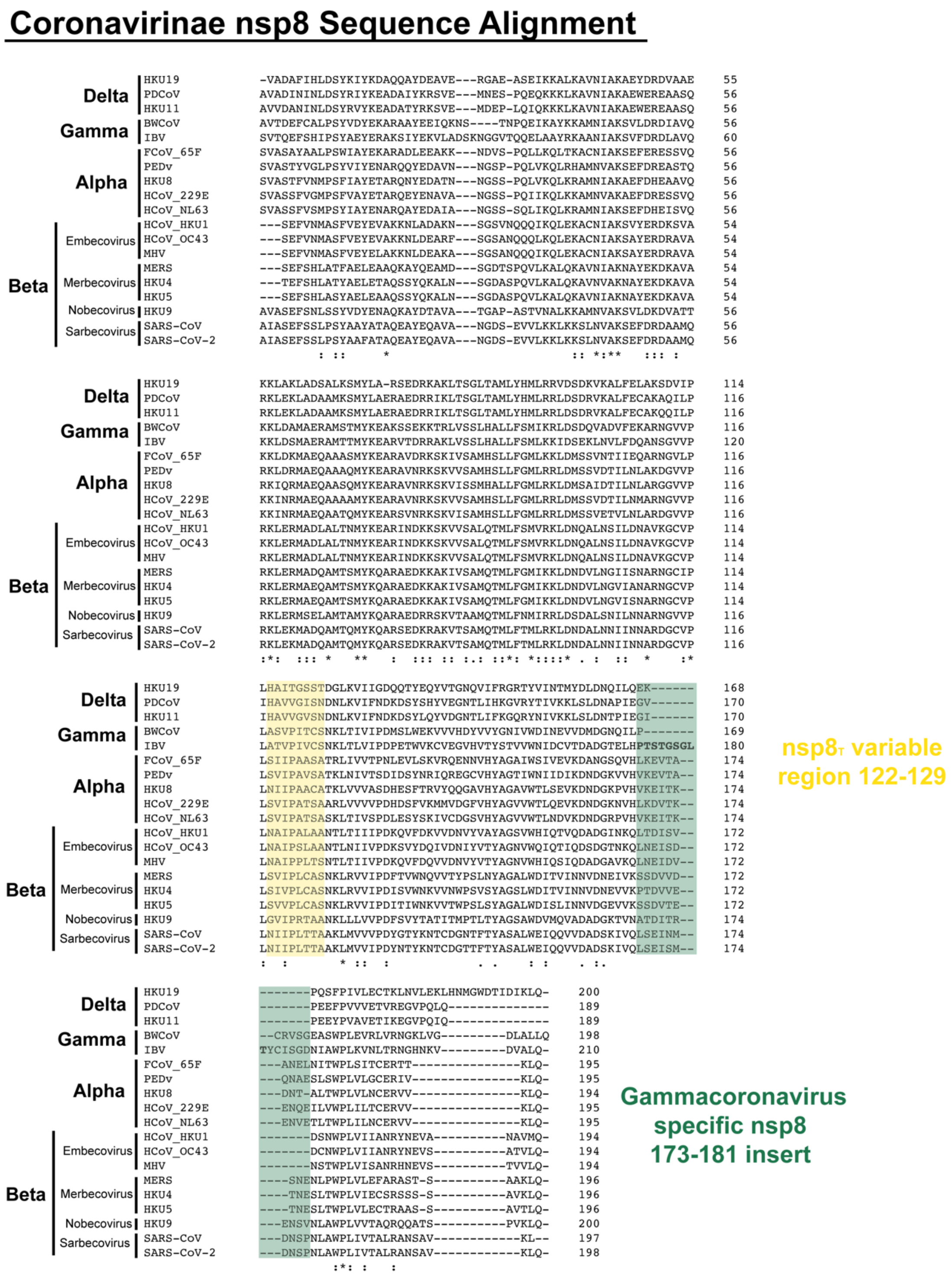


**Figure S6: Coronavirinae nsp8 sequence alignment.** Clustal omega alignment of coronavirus nsp8 proteins from alpha, beta, gamma, and delta genera. "*" notation means residue is fully conserved, ":" means residue is strongly conserved, and "." Denotes a weakly conserved residue.

#### Gammacoronavirus genus nsp8 sequence alignment

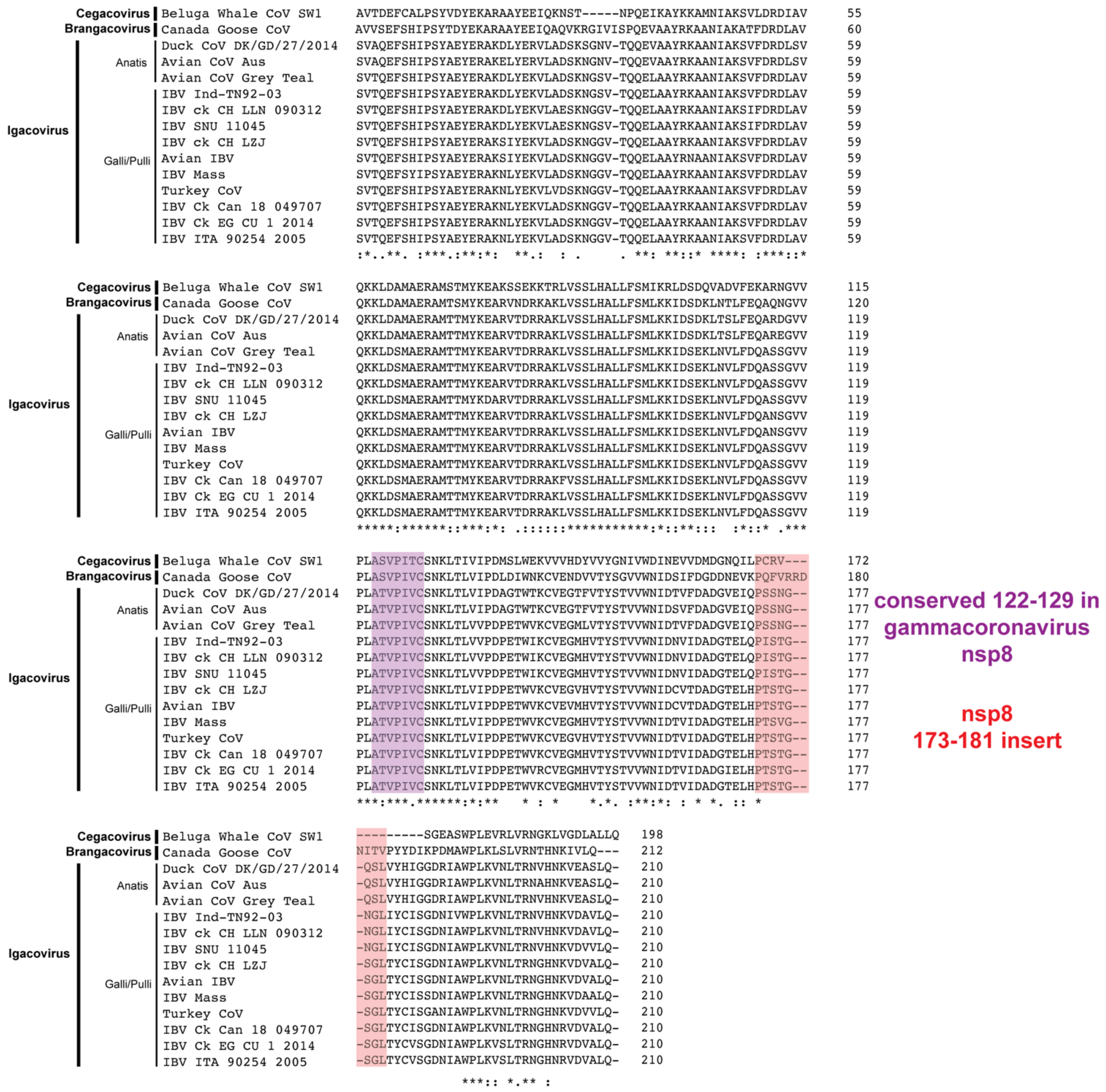


**Figure S7 Gammacoronavirus nsp8 sequence alignment:** Clustal omega alignment of nsp8 proteins from viruses residing within the gammacoronavirus genus. “*” notation means residue is fully conserved, “:” means residue is strongly conserved, and “.” Denotes a weakly conserved residue.

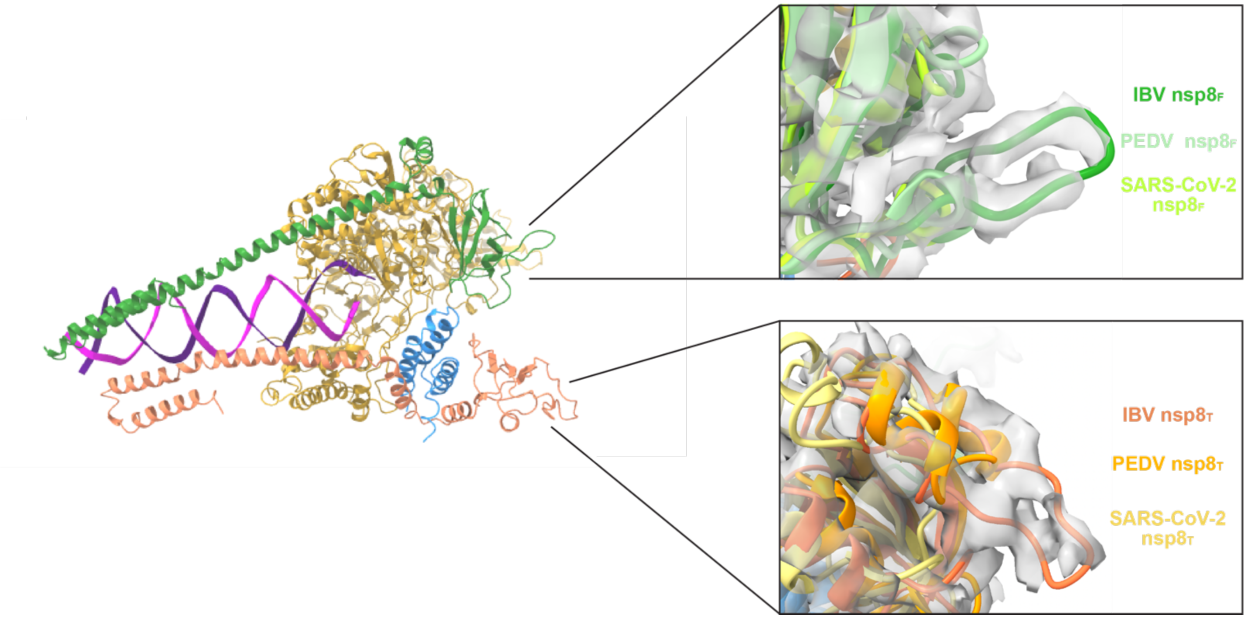


**Figure S8 Structural view of IBV specific nsp8 173-181 insert:** Cartoon model of the IBV complex (left) with zoomed in views of superimposed models of IBV, PEDv (8URB) and SARS-CoV-2 (6XEZ). Focused view highlights IBV nsp8 insert in both protomers found on RTC model with supporting electron density.

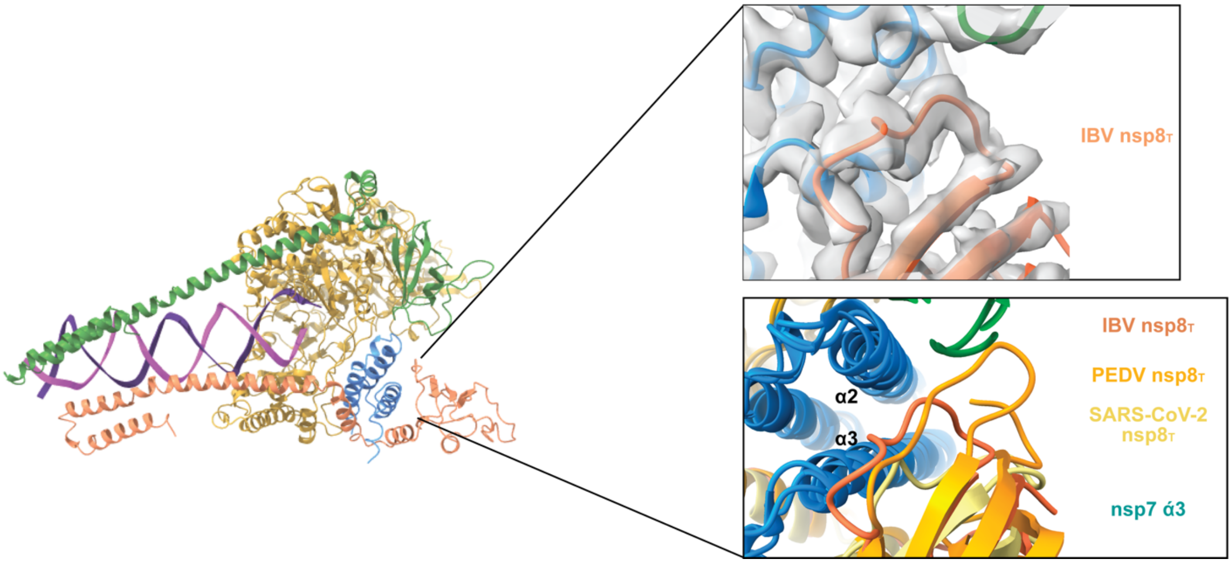


**Figure S9 Structural view of nsp8$_T$ 122-129 variable region:** Cartoon model of the IBV complex (left) with zoomed in views of superimposed models of IBV, PEDv (8URB) and SARS-CoV-2 (6XEZ). Focused view highlights IBV nsp8$_T$ 122-129 that has observed variability across genera. IBV model electron density shown to support conformation.

### Coronavirinae nsp12 Sequence Alignment

```
                   HKU19       ---NSYLNRVKGSS-DARLEPCTSDNRPDVVVRAFNIYNN--ATAGIFKSTKNNCTRFKS   54
Delta              PDCoV       ---SAYLNRVTGSS-DARLEPLQPGTQPDAVKRAFHVHND--TTSGIFLSTKSNCARFKT   54
                   HKU11       ---SPYLNRVTGSS-GARLEPQQPGVTPDAVKRAFHVHNN--TTSGIFLSTKTNCSRFRT   54
                   FCOV_65F    ---QELFKRVRGSS-AARLEPC-NGTDPDHVSRAFDIYNK--DVACIGKFLKTNCSRFRN   53
                   HCOV_229E   ---SSYLNRVRGSS-AARLEPC-NGTDIDYCVRAFDVYNK--DASFIGKNLKSNCVRFKN   53
Alpha              PEDv        ----GLFKRVRGSS-AARLEPC-NGTDTQHVYRAFDIYNK--DVACLGKFLKVNCVRLKN   52
                   HKU8        ---NNYLNRVRGSS-AARLEPC-NGTEPEHVIRAFDIYNK--DVACIGKFVKVNCVRFKN   53
                   HCOV_NL63   ----SYLNRARGSS-AARLEPC-NGTDIDKCVRAFDIYNK--NVSFLGKCLKMNCVRFKN   52
Gamma              BWCoV       ---QNYLNRVRGLS-EARLRPCASGLLPDVVKRAFDLYNS--NTAGMYASLKHNCARFQE   54
                   IBV         ---KNYLNRVRGSS-EARLIPLANGCDPDVVKRAFDVCNK--ESAGMFQNLKRNCARFQE   54
                   HCoV_OC43   ---TNFLNRVRGASVDARLVPCANGLSTDVQLRAFDIYNA--SVAGIGLHLKVNCCRFQR   55
Embecovirus        HCoV_HKU1   ---LNFLNRVRGTSVNARLVPCASGLSTDVQLRAFDICNT--NRAGIGLYYKVNCCRFQR   55
                   MHV         -----FFKRVRGTSVNARLVPCASGLDTDVQLRAFDICNA--NRAGIGLYYKVNCCRFQR   53
                   MERS        ---SNFLNRVRGSIVNARIEPCSSGLSTDVVFRAFDICNYKAKVAGIGKYYKTNTCRFVE   57
Beta Merbecovirus  HKU4        ---TNFLNRVRGSSVNARLEPCSSGLTTDVVYRAFDICNFKARVAGIGKYYKTNTCRFVQ   57
                   HKU5        ---SNFLNRVRGSIVNARIEPCASGLTTDVVFRAFDICNYKAKVAGIGKYYKTNTCRFVE   57
Nobecovirus        HKU9        ARDECFLNRVRGTSGVARLVPLGSGVQPDIVLRAFDICNT--KVAGFGLHLKNNCCRYQE   58
Sarbecovirus       SARS-CoV    ----TFLNRVCGVS-AARLTPCGTGTSTDVVYRAFDIYNE--KVAGFAKFLKTNCCRFQE   53
                   SARS-CoV-2  ---QSFLNRVCGVS-AARLTPCGTGTSTDVVYRAFDIYND--KVAGFAKFLKTNCCRFQE   54
                                     ::*. *    **: *   .   :   ***.: *     : :    * *  *

                   HKU19       TRPGSIINKPVRTIETFFVTKQCTENVFRAEEQCYNMLPKSIVSTDDKFSCVAYHDFFKF   114
Delta              PDCoV       TRSALPLP-NKGEVELYFVTKQCAAKVFEIEEECYNALSTELYTTDDTFGVLAKTEFFKF   113
                   HKU11       TKQNLPLP-NKGSVELYFVSKQCSQQVFEIEETCYNMFDDSLKSTPEKFGVLARTEFFKF   113
                   FCoV_65F    LD----------KHDAYYVVKRCTKNVMDHEQVCYNDLKD--------SGAVAEHDFFLY   95
                   HCoV_229E   VD----------KDDAFYIVKRCIKSVMDHEQSMYNLLKG--------CNAVAKHDFFTW   95
Alpha              PEDv        LD----------KHDAFYVVKRCTKSAMEHEQSIYSRLEK--------CGAVAEHDFFTW   94
                   HKU8        AD----------KHDAFYVVKRCTKSVMEHEQSIYDALKD--------CGAVSPHDFFVW   95
                   HCoV_NL63   AD----------LKDGYFVIKRCTKSVMEHEQSMYNLLNF--------SGALAEHDFFTW   94
Gamma              BWCoV       LDENDD------EIDSFFVVKQTTPHNFEHEEKCYLDLKA---------DCVAVHDFFRF   99
                   IBV         VRDTED--GNLEYCDSYFVVKQTTPSNYEHEKACYEDLKS---------EVTADHDFFVF   103
                   HCoV_OC43   VDENGD------KLDQFFVVKRTDLTIYNREMKCYERVKD--------CKFVAEHDFFTF   101
Embecovirus        HCoV_HKU1   IDDDGN------KLDKFFVVKRTNLEVYNKEKTYYELTKS--------CGVVAEHDFFTF   101
                   MHV         VDEDGN------KLDKFFVVKRTNLEVYNKEKECYELTKD--------CGVVAEHEFFTF   99
                   MERS        LDDQGH------HLDSYFVVKRHTMENYELEKHCYDLLRD--------CDAVAPHDFFIF   103
Beta Merbecovirus  HKU4        VDDEGH------KLDSYFIVKRHTMSNYELEKRCYDLLKD--------CDAVAIHDFFIF   103
                   HKU5        VDDEGH------RLDSFFVVKRHTMENYELEKRCYDLVKD--------CDAVAVHDFFIF   103
Nobecovirus        HKU9        LDADGT------QLDSYFVVKRHTESNYLLEQRCYEKLKD--------CGVVARHDFFKF   104
Sarbecovirus       SARS-CoV    KDEEGN------LLDSYFVVKRHTMSNYQHEETIYNLVKD--------CPAVAVHDFFKF   99
                   SARS-CoV-2  KDEDDN------LIDSYFVVKRHTFSNYQHEETIYNLLKD--------CPAVAKHDFFKF   100
                                             : ::: *:        *   *                 :  :** :

                   HKU19       DG---VNNVVRRHLTKYTLLDLVYALRHLSTS-QEIIQEILTTMCGTSEE---WF--VD   164
Delta              PDCoV       DK---IPNVNRQYLTKYTLLDLAYALRHLSTS-KDVIQEILITMCGTPED---WF--GE   163
                   HKU11       DK---IPNVNRQFLTKYTLLDLAYALRHLSTS-RDVIKEILITICGTTEE---WF--GD   163
                   FCoV_65F    KEGRCEFGNVARKDLTKYTMMDLCYAIRNFDEKNCEVLKEILVTLGACNES----FFENK   151
                   HCoV_229E   HEGRTIYGNVSRQDLTKYTMMDLCFALRNFDEKDCGVFKEILVLTGCCNTD----YFEMK   151
Alpha              PEDv        KDGRAIYGNVCRKDLTEYTMMDLCYALRNFNENNCDVLKSILIKVGACEES----YFNNK   150
                   HKU8        KDGRSVYGNIARHDLTKYTMMDLVHALRNFDEKNCETLKEILVISGACDSS----YFDNK   151
                   HCoV_NL63   KDGRVIYGNVSRHNLTKYTMMDLVYAMRNFDEQNCDVLKEVLVLTGCCDNS----YFDSK   150
Gamma              BWCoV       E-G---MYSICRQRLTKYTMMDLCYAFRHFDPNDCDVLKEILVVKGCCEWD----YFDQP   151
                   IBV         NKN---IYNISRQRLTKYTMMDFCYALRHFDPKDCEVLKEILVTYGCIEDYHPKWFEENK   160
                   HCoV_OC43   DVEGSRVPHIVRKDLTKYTMLDLCYALRHFDRNDCMLLCDILSIYAGCEQS----YFTKK   157
Embecovirus        HCoV_HKU1   DIDGSRVPHIVRKNLSKYTMLDLCYALRHFDCNDCSVLCEILCEYADCKES----YFSKK   157
                   MHV         DVEGSRVPHIVRKDLSKFTMLDLCYALRHFDRNDCSTLKEILLTYAECDES----YFQKK   155
                   MERS        DVDKVKTPHIVRQRLTEYTMMDLVYALRHFDQN-SEVLKAILVKYGCCDVT----YFENK   158
Beta Merbecovirus  HKU4        DVDKTKTPHIVRQSLTEYTMMDLVYALRHFDQNNCEVLKSILVKYGCCEQS----YFDNK   159
                   HKU5        DVDKVKTPHIVRQRLTEYTMMDLVYALRHFDQNNCEVLKSILVKYGCCDAS----YFDNK   159
Nobecovirus        HKU9        NIEGVMTPHVSRERLTKYTMADLVYSLRHFDNNNCDTLKEILVLRGCCTAD----YFDRK   160
Sarbecovirus       SARS-CoV    RVDGDMVPHISRQRLTKYTMADLVYALRHFDEGNCDTLKEILVTYNCCDDD----YFNKK   155
                   SARS-CoV-2  RIDGDMVPHISRQRLTKYTMADLVYALRHFDEGNCDTLKEILVTYNCCDDD----YFNKK   156
                                        : *. *:::*: *: .::*::.      :  :*             :

                   HKU19       GWYDPIENPTFYDEFHKLGSLINNCVVMANKFADTCKTVGLVGILTADNQDLGGQIYDFG   224
Delta              PDCoV       NWFDPIENPSFYKEFHKLGDILNRCVLNANKFASACIDAGLVGILTPDNQDLLGQIYDFG   223
                   HKU11       SWFDPIENPTFYREFHKLGSVLNRCVLNANAFAKACSELGIVGILTPDNQDLLGQIYDFG   223
                   FCoV_65F    DWFDPVENEAIHEVYARLGPIVANAMLKCVAFCDAIVAKGYIGIITLDNQDLNGNFYDFG   211
                   HCoV_229E   NWFDPIENEDIHRVYAALGKVVANAMLKCVAFCDEMVLKGVVGVLTLDNQDLNGNFYDFG   211
Alpha              PEDv        VWFDPVENEDIHRVYAVLGTIVSRAMLKCVKFCDAMVEQGIVGVVTLDNQDLNGDFYDFG   210
                   HKU8        NWYDPVENEDIHRVYAKLGCVVANAMLKCVALCDAMVAKGVVGVLTLDNQDLNGNFYDFG   211
                   HCoV_NL63   GWYDPVENEDIHRVYASLGKIVARAMLKCVALCDAMVAKGVVGVLTLDNQDLNGNFYDFG   210
Gamma              BWCoV       NWYDPVENPDWFSLISRLGPIFQRALIKVAEFCDLMVEKGYIGVVTLDNQDLNGNFYDFG   211
                   IBV         DWYDPIENPKYYAMLAKMGPIVRRALLNAIEFGNLMVEKGYVGVITLDNQDLNGKFYDFG   220
                   HCoV_OC43   DWYDFVENPDIINVYKKLGPIFNRALVSATEFADKLVEVGLVGVLTLDNQDLNGKWYDFG   217
Embecovirus        HCoV_HKU1   DWYDFVENPDIINIYKKLGPIFNRALLNTVSFADTLVKVGLVGVLTLDNQDLYGQWYDFG   217
                   MHV         DWYDFVENPDIINVYKKLGPIFNRALLNTAKFADTLVEAGLVGVLTLDNQDLYGQWYDFG   215
                   MERS        LWFDFVENPSVIGVYHKLGERVRQAILNTVKFCDHMVKAGLVGVLTLDNQDLNGKWYDFG   218
Beta Merbecovirus  HKU4        LWFDFVENPSVIGVYHKLGERIRQAMLNTVKMCDHMVKSGLVGVLTLDNQDLNGKWYDFG   219
                   HKU5        LWFDFVENPNVISVYHKLGERIRQAVLNTVKFCDQMVKSGLVGVLTLDNQDLNGKWYDFG   219
Nobecovirus        HKU9        DWYDPVENPDIIRVYHNLGETVRKAVLSAVKMADSMVEQGLIGVLTLDNQDLNGQWYDFG   220
Sarbecovirus       SARS-CoV    DWYDFVENPDILRVYANLGERVRQALLKTVQFCDAMRDAGIVGVLTLDNQDLNGNWYDFG   215
                   SARS-CoV-2  DWYDFVENPDILRVYANLGERVRQALLKTVQFCDAMRNAGIVGVLTLDNQDLNGNWYDFG   216
                                *:* :**         :*  . ..::    : .     * :*::* ***** *. ****
```

IBV nsp12
67-72 insert

IBV nsp12
113-115
shortened loop

IBV nsp12
156-169 loop
insert

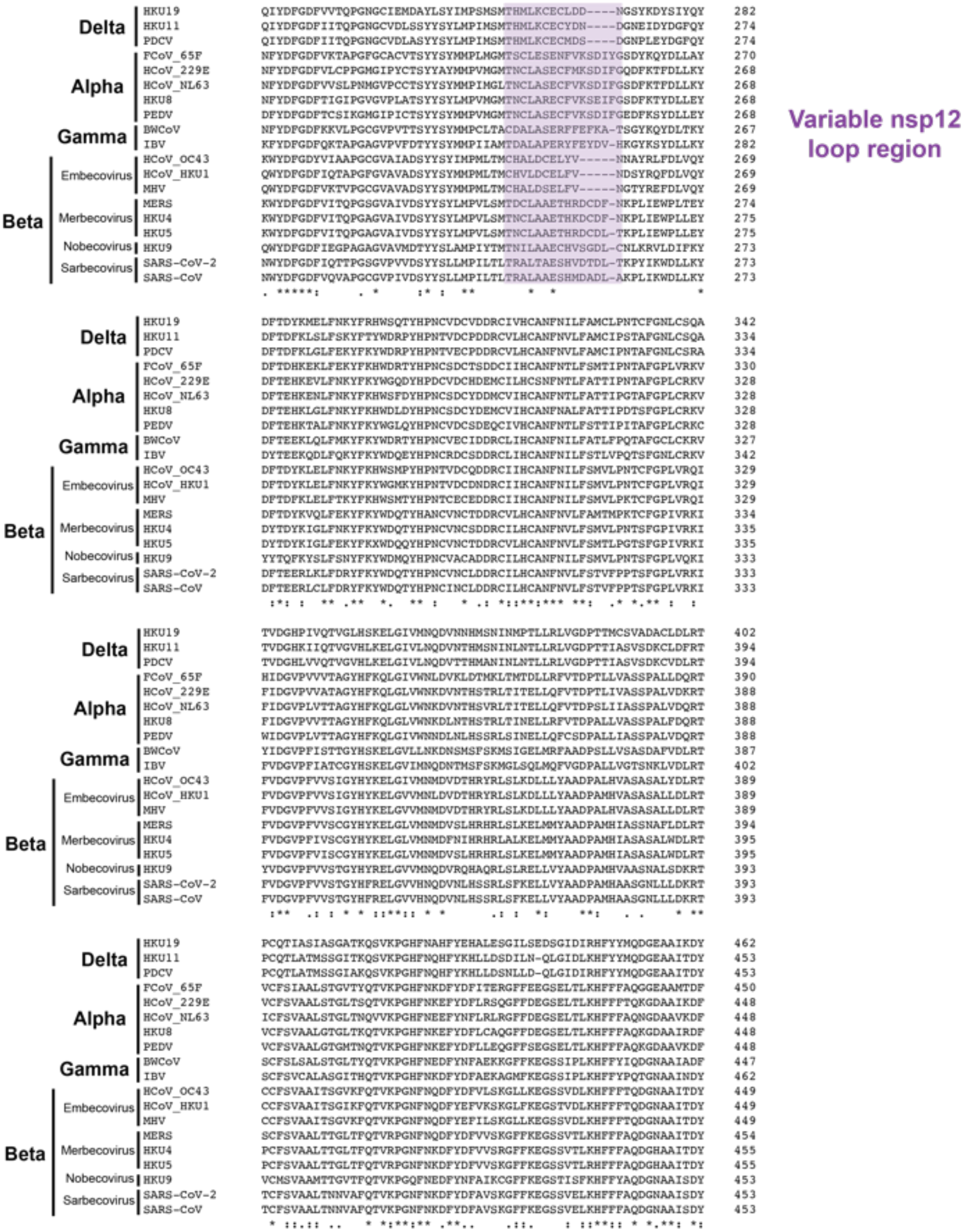
Variable nsp12 loop region
Delta
Alpha
Gamma
Beta
Embecovirus
Merbecovirus
Nobecovirus
Sarbecovirus
HKU19
HKU11
PDCV
FCoV_65F
HCoV_229E
HCoV_NL63
HKU8
PEDV
BWCoV
IBV
HCoV_OC43
HCoV_HKU1
MHV
MERS
HKU4
HKU5
HKU9
SARS-CoV-2
SARS-CoV

```
Delta        HKU19        SYYRYNTPTMVDIKQFLFVMEVADKYLSFYDGGCIPAETVVVNNLDKSAGYPFNKLGKAR   522
             HKU11        SYYRYNTPTMVDIKMFLFVLEVADKYLQPYEGGCLNAQSVVVNNLDKYAGYPFNKLGKAR   513
             PDCV         SYYRYNTPTMVDIKMFLFCLEVADKYLEPYEGGCINAQSVVVSNLDKSAGYPFNKLGKAR   513
Alpha        FCoV_65F     NYYRYNRVTVLDICQAQFVYKIVCKYFDCYDGGCINAREVVVTNYDKSAGYPLNKFGKAR   510
             HCoV_229E    DYYRYNRPTILDIGQARVAYQVAARYFDCYEGGCITSREVVVTNLNKSAGWPLNKFGKAG   508
             HCoV_NL63    DFYRYNKPTILDICQARVTYKIVSRYFDIYEGGCIKACEVVVTNLNKSAGWPLNKFGKAS   508
             HKU8         DFYRYNRPTVLDICQARVAYHVVKRYFDIYEGGCIAARDVVVTNLNKSAGYPLNKFGKAS   508
             PEDV         DYYRYNRPTVLDICQARVVYQIVQRYFDIYEGGCITAKEVVVTNLNKSAGYPLNKFGKAG   508
Gamma        BWCoV        DYYRFNKPTMVDIQQFLFCFEVTDKYFEQYDGGCIPANQVVVTNLDKSAGYPFNKFGKAR   507
             IBV          DYYRYNRPTMFDIRQLLFCLEVTSKYFECYEGGCIPASQVVVNNLDKSAGYPFNKFGKAR   522
Beta  Embecovirus   HCoV_OC43  NYYKYNLPTMVDIKQLLFVLEVVYKYFEIYDGGCIPASQVIVNNYDKSAGYPFNKFGKAR   509
                     HCoV_HKU1  NYYKYNLPTMVDIKQLLFVLEVVYKYFEIYDGGCIPASQVIVNNYDKSAGYPFNKFGKAR   509
                     MHV        NYYKYNLPTMVDIKQLLFVLEVVNKYFEIYEGGCIPATQVIVNNYDKSAGYPFNKFGKAR   509
      Merbecovirus   MERS       NYYSYNLPTMCDIKQMLFCMEVVNKYFEIYDGGCLNASEVVVNNLDKSAGHPFNKFGKAR   514
                     HKU4       SYYAYNLPTMVDIKQMLFCMEVVDKYFDIYDGGCLNASEVIVNNLDKSAGHPFNKFGKAR   515
                     HKU5       SYYAYNLPTMCDIKQMLFCMEVVDRYFEIYDGGCLNASEVIVNNLDKSAGHPFNKFGKAR   515
      Nobecovirus    HKU9       DYYRYNLPTMCDIKQLLFSLEVVDKYFDCYDGGCLQASQVVVANYDKSAGFPFNKFGKAR   513
      Sarbecovirus   SARS-CoV-2 DYYRYNLPTMCDIRQLLFVVEVVDKYFDCYDGGCINANQVIVNNLDKSAGFPFNKWGKAR   513
                     SARS-CoV   DYYRYNLPTMCDIRQLLFVVEVVDKYFDCYDGGCINANQVIVNNLDKSAGFPFNKWGKAR   513
                                .:* :*  *: **    .  .:. :*:. *:***: :  *:* * :* ** *:** ***

Delta        HKU19        NYYD-LTYAEQNAMFEYTKRNVLPTVTQMNLKYAISAKDRARTVAGVSIISTMTNRQYHQ   581
             HKU11        NYYD-MTYAEQNQLFEYTKRNVLPTLTQMNLKYAISAKDRARTVAGVSIISTMTNRQYHQ   572
             PDCV         NYYD-MTHAEQNQLFEYTKRNVLPTLTQMNLKYAISAKDRARTVAGVSIISTMTNRQYHQ   572
Alpha        FCoV_65F     LYYETLSYEEQDAIFALTKRNVLPTMTQMNLKYAISGKARARTVGGVSLLSTMTTRQYHQ   570
             HCoV_229E    LYYESISYEEQDAIFLLTKRNILPTMTQLNLKYAISGKERARTVGGVSLLATMTTRQFHQ   568
             HCoV_NL63    LYYESISYEEQDALFALTKRNVLPTMTQLNLKYAISGKERARTVGGVSLLSTMTTRQYHQ   568
             HKU8         LYYESLSYEEQDALYALTKRNVLPTMTQLNLKYAISGKERARTVGGVSLLSTMTTRQFHQ   568
             PEDV         LYYESLSYEEQDELYAYTKRNILPTMTQLNLKYAISGKERARTVGGVSLLSTMTTRQYHQ   568
Gamma        BWCoV        LYYESLSYAEQDQLFELTKRNILPTITQINMKYAISAKSRARTVAGVSILSSMTNRQFHQ   567
             IBV          LYYE-MSLEEQDQLFESTKKNVLPTITQMNLKYAISAKNRARTVAGVSILSTMTNRQFHQ   581
Beta  Embecovirus   HCoV_OC43  LYYEALSFEEQDEIYAYTKRNVLPTLTQMNLKYAISAKNRARTVAGVSILSTMTGRMFHQ   569
                     HCoV_HKU1  LYYEALSFEEQNEIYAYTKRNVLPTLTQMNLKYAISAKNRARTVAGVSILSTMTGRMFHQ   569
                     MHV        LYYEALSFEEQDEIYAYTKRNVLPTLTQMNLKYAISAKNRARTVAGVSILSTMTGRMFHQ   569
      Merbecovirus   MERS       VYYESMSYQEQDELFAMTKRNVIPTMTQMNLKYAISAKNRARTVAGVSILSTMTNRQYHQ   574
                     HKU4       VYYESMSYQEQDELFAVTKRNVLPTITQMNLKYAISAKNRARTVAGVSILSTMTNRQYHQ   575
                     HKU5       VYYESLSYQEQDELFAMTKRNVLPTITQMNLKYAISAKNRARTVAGVSILSTMTNRQYHQ   575
      Nobecovirus    HKU9       LYYESLSYADQDELFAYTKRNVLPTITQMNLKYAISAKNRARTVAGVSIASTMTNRQFHQ   573
      Sarbecovirus   SARS-CoV-2 LYYDSMSYEDQDALFAYTKRNVIPTITQMNLKYAISAKNRARTVAGVSICSTMTNRQFHQ   573
                     SARS-CoV   LYYDSMSYEDQDALFAYTKRNVIPTITQMNLKYAISAKNRARTVAGVSICSTMTNRQFHQ   573
                                 **: ::  :*: ::  **:*::**:**:*:*****.* *****.***: ::** * :**

Delta        HKU19        KLLKSISVARNQTIVIGTTKFYGGWDNMLRNLMANINNPKLAGWDYPKCDRSMPNLLRIT   641
             HKU11        KMLKSISLARNQTIVIGTTKFYGGWDNMLRRLMNGINNPILVGWDYPKCDRSMPNMLRIA   632
             PDCV         KMLKSISLARNQTIVIGTTKFYGGWDNMLRRLMCNINNPILVGWDYPKCDRSMPNMLRIA   632
Alpha        FCoV_65F     KHLKSIAATRNATVVIGTTKFYGGWDNMLKNLMRDVDNGCLMGWDYPKCDRALPNMIRMA   630
             HCoV_229E    KCLKSIVATRNATVVIGTTKFYGGWDNMLKNLMADVDDPKLMGWDYPKCDRAMPSMIRML   628
             HCoV_NL63    KHLKSIVNTRNATVVIGTTKFYGGWNNMLRTLIDGVENPMLMGWDYPKCDRALPNMIRMI   628
             HKU8         KHLKSIVNTRNATVVIGTTKFYGGWDNMLRNLMDGVDNACLMGWDYPKCDRALPNMIRMI   628
             PEDV         KHLKSIVNTRGASVVIGTTKFYGGWDNMLKNLIDGVENPCLMGWDYPKCDRALPNMIRMI   628
Gamma        BWCoV        KCLKSIVNTRNATVVIGTTKFYGGWDNMLRNLMRGVEDPVLMGWDYPKCDRAVPSLLRLS   627
             IBV          KILKSIVNTRNAPVVIGTTKFYGGWDNMLRNLIQGVEDPILMGWDYPKCDRAMPNLLRIA   641
Beta  Embecovirus   HCoV_OC43  KCLKSIAATRGVPVVIGTTKFYGGWDDMLRRLIKDVDNPVLMGWDYPKCDRAMPNLLRIV   629
                     HCoV_HKU1  KCLKSIAATRGVPVVIGTTKFYGGWDDMLRHLIKDVDNPVLMGWDYPKCDRAMPNILRIV   629
                     MHV        KCLKSIAATRGVPVVIGTTKFYGGWDDMLRRLIKDVDSPVLMGWDYPKCDRAMPNILRIV   629
      Merbecovirus   MERS       KMLKSMAATRGATCVIGTTKFYGGWDFMLKTLYKDVDNPHLMGWDYPKCDRAMPNMCRIF   634
                     HKU4       KMLKSMAATRGATCVIGTTKFYGGWDFMLKTLYKDVESPHLMGWDYPKCDRAMPNMCRIL   635
                     HKU5       KMLKSMAATRGSTCVIGTTKFYGGWDFMLKTLYKDVDNPHLMGWDYPKCDRAMPNMCRIF   635
      Nobecovirus    HKU9       KMLKSIAAARGASVVIGTTKFYGGWNRMLRTLCEGVDNPHLMGWDYPKCDRAMPNLLRIF   633
      Sarbecovirus   SARS-CoV-2 KLLKSIAATRGATVVIGTSKFYGGWHNMLKTVYSDVENPHLMGWDYPKCDRAMPNMLRIM   633
                     SARS-CoV   KLLKSIAATRGATVVIGTSKFYGGWHNMLKTVYSDVETPHLMGWDYPKCDRAMPNMLRIM   633
                                * ***:  :*.   ****:******. **: :  .::   * *********::*.: *:

Delta        HKU19        SSLLLARKH-ACCTHSQRFYRLANECAQVLSEIVVSGNVMYVKPGGTSSGDATTAYANSV   700
             HKU11        ASCLLARKH-TCCNQSQRFYRLANECCQVLSEVVVSGNNLYVKPGGTSSGDATTAYANSV   691
             PDCV         ASCLLARKH-TCCNQSQRFYRLANECCQVLSEVVVSGNNLYVKPGGTSSGDATTAYANSV   691
Alpha        FCoV_65F     SAMVLGSKHIGCCTHSDRFYRLSNELAQVLTEVVHCTGGFYIKPGGTTSGDGTTAYANSA   690
             HCoV_229E    SAMILGSKHVTCCTASDKFYRLSNELAQVLTEVVYSNGGFYFKPGGTTSGDATTAYANSV   688
             HCoV_NL63    SAMVLGSKHVNCCTATDRFYRLGNELAQVLTEVVYSNGGFYFKPGGTTSGDASTAYANSI   688
             HKU8         SAMILGSKHVNCCTNSDRYYRLCNELAQVLTEVVYSNGGFYMKPGGTTSGDATTAYANSV   688
             PEDV         SAMILGSKHTTCCSSTDRFFRLCNELAQVLTEVVYSNGGFYLKPGGTTSGDATTAYANSV   688
Gamma        BWCoV        ASLILARRHKGCCDWNERIYRLANEAAQVLSEVALSNGGLYVKPGGTSSGDATTAYANSA   687
             IBV          ASLVLARKHTNCCTWSERVYRLYNECAQVLSETVLATGGIYVKPGGTSSGDATTAYANSV   701
Beta  Embecovirus   HCoV_OC43  SSLVLARKHETCCSQSDRFYRLANECAQVLSEIVMCGGCYYVKPGGTSSGDATTAFANSV   689
                     HCoV_HKU1  SSLVLARKHEFCCSHGDRFYRLANECAQVLSEIVMCGGCYYVKPGGTSSGDATTAFANSV   689
                     MHV        SSLVLARKHDSCCSHTDRFYRLANECAQVLSEIVMCGGCYYVKPGGTSSGDATTAFANSV   689
      Merbecovirus   MERS       ASLILARKHGTCCTTRDRFYRLANECAQVLSEYVLCGGGYYVKPGGTSSGDATTAYANSV   694
                     HKU4       ASLILARKHSTCCTNSDRFYRLANECAQVLSEYVLCGGGYYVKPGGTSSGDATTAYANSV   695
                     HKU5       ASLILARKHSTCCTNTDRFYRLANECAQVLSEYVLCGGGYYVKPGGTSSGDATTAYANSV   695
      Nobecovirus    HKU9       ASLILARKHSTCCNASERFYRLANECAQVLSEMVLCGGGFYVKPGGTSSGDSTTAYANSV   693
      Sarbecovirus   SARS-CoV-2 ASLVLARKHTTCCSLSHRFYRLANECAQVLSEMVMCGGSLYVKPGGTSSGDATTAYANSV   693
                     SARS-CoV   ASLVLARKHNTCCNLSHRFYRLANECAQVLSEMVMCGGSLYVKPGGTSSGDATTAYANSV   693
                                :: :*. :*  **   .: :** ** .***:* . . .  *.*****:***.:**:***
```

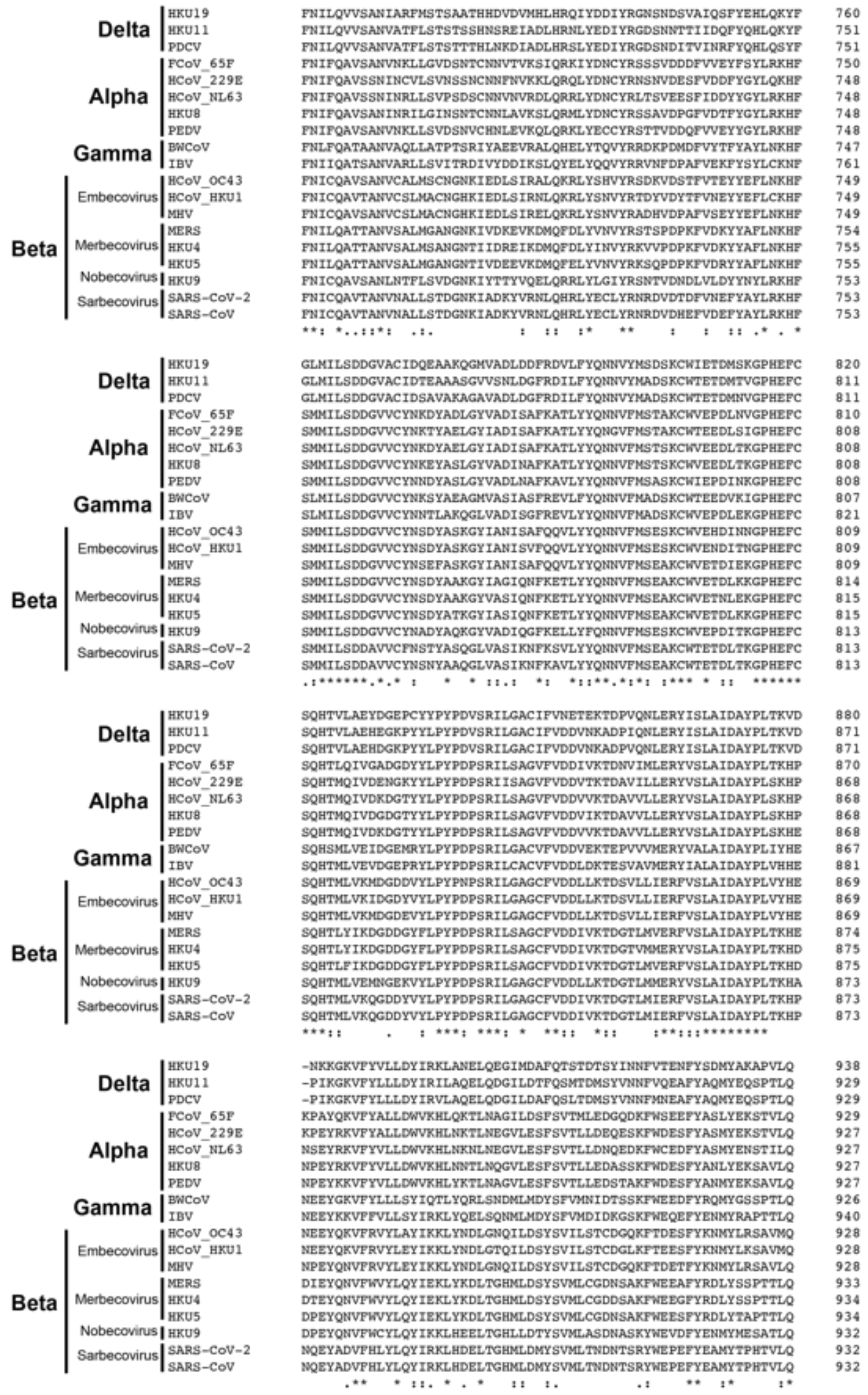


**Figure S10 Coronavirinae nsp12 sequence alignment:** Clustal omega alignment of coronavirus nsp12 proteins from alpha, beta, gamma, and delta genera. "*" notation means residue is fully conserved, ":" means residue is strongly conserved, and "." Denotes a weakly conserved residue.

### Gammacoronavirus genus nsp12 sequence alignment

```
Cegacovirus    | Beluga Whale CoV SW1     ----------SVPQNYLNRVRGLSEARLRPCASGLLPDVVKRAFDLYNSNTAGMYASLKH   50
Brangacovirus  | Canada Goose CoV         -------------NNYLNRVRGSSEARLIPLTDGVQPDVVKRAFDICNKDAAGMFQNLKH   47
               | Duck CoV DK/GD/27/2014   ----SDAGAPGFDKNYLNRVRGSSEARLIPLSDGNQPHIVKRAFDVCNKESAGMFQNMKH   56
        Anatis | Avian CoV Aus            STYVAPVVSTNSDKNYLNRVRGSSEARLIPLSDGNQPHIVKRAFDVCNKESAGMFQNMKH   60
               | Avian CoV Grey Teal      GTYVAPTVSPSVDKNYLNRVRGSSEARLIPLSDGNQPHIVKRAFDVCNKESAGMFQNMKH   60
               | IBV Ind-TN92-03          ----SVAVASDFDKNYLNQGTGWQCGSAVTPSSWMWPDVVWRAFDVCNKESAGMFQNLKR   56
               | IBV ck CH LLN 090312     ----SDAGAPGFDKNYLNRVRGSSEARLIPLANGCEPDVVERAFDVCNKESAGMFKNLKR   56
Igacovirus     | IBV SNU 11045            ----SDVGVPGFDKNYLNRVRGSSEARLIPLANGCEPDVVERAFDVCNKESAGMFKNLKR   56
               | IBV ck CH LZJ            ----SDAGAPGFDKNYLNRVRGSSEARLIPLANGCEPDVVERAFDVCNKESAGMFKNLKR   56
   Galli/Pulli | Avian IBV                ----SVAGASDFDKNYLNRVRGSSEARLIPLASGCDPDVVKRAFDVCNKESAGMFQNLKR   56
               | IBV Mass                 ----SVAVASGFDKNYLNRVRGSSEARLIPLANGCDPDVVKRAFDVCNKESAGMFQNLKR   56
               | Turkey CoV               ----SVAGASDFGKNYLNRVRGSSEARLIPLASGCDPDVVKRAFDVCNKESAGMFINFKR   56
               | IBV Ck Can 18 049707     ----SAAGAPDFDKNYLNRVRGSSEARLIPLANGCDPDVVMRAFDVCNKESAGMFKNFKR   56
               | IBV Ck EG CU 1 2014      ----SAAGVADFDKNYLNRVRGSSEARLIPLSIGCDPDVVKRAFDVCNKESAGMFQNLKR   56
               | IBV ITA 90254 2005       ----SAAGVADSDKNYLNRVRGSSEARLIPLSSGCDPDVVKRAFDVCNKESAGMFRNLKR   56
                                                       :****:  * . .     :    *.:* ****: *.::***: .:*:
```

```
Cegacovirus    | Beluga Whale CoV SW1     NCARFQELDENDD-------EIDSFFVVKQTTPHNFEHEEKCYLDLKADCVAVHDFFRFE  103
Brangacovirus  | Canada Goose CoV         NCARFQEL--GPEDYKKGFPYLNSFFVVKQTTPANYQHELSVYQDLKSEVVAVHDFFIFD  105
               | Duck CoV DK/GD/27/2014   NCARFQEVFEGEDSWKRGLEYKDSYFVVKQTTPSNYAHEQACYEDLKSDVVAVHDFFVFN  116
        Anatis | Avian CoV Aus            NCARFQEVVEGEDSWKHGLEYKDSYFVVKQTTPSNYVHEQACYEDLKSDVVAVHDFFVFN  120
               | Avian CoV Grey Teal      NCARFQEVFESEDSWKRGLEYKDSYFVVKQTTPSNYAHEQACYEDLKSDVVAVHDFFVFN  120
               | IBV Ind-TN92-03          KCGRFQEVRDTED---GILEYCDSYFGVKQTTPSNYEHEKACYEELKSEVTADHDFFVFN  113
               | IBV ck CH LLN 090312     NCARFQEVCGTED---GNLEYRDSYFVVKQTTPTNYEHEKACYEDLKSEVTADHDFFVFN  113
Igacovirus     | IBV SNU 11045            NCARFQEVCGTED---GNLEYRDSYFVVKQTTPSNYEHEKACYEDLKSEVTADHDFFVFN  113
               | IBV ck CH LZJ            NCARFQEVCGTED---GNLEYRDSYFVVKQTTPSNYEHEKACYEDLKSEVTADHDFFVFN  113
   Galli/Pulli | Avian IBV                NCARFQELRDTED---GNLEYLDSYFVVKQTTPSNYEHEKSCYEDLKSEVTADHDFFVFN  113
               | IBV Mass                 NCARFQEVRDTED---GNLEYCDSYFVVKQTTPSNYEHEKACYEDLKSEVTADHDFFVFN  113
               | Turkey CoV               NCARFQEVRDTED---GNLEYLNSYFVVKQTTPSNYEHEKSCYEDLKSEVTADHDFFVFN  113
               | IBV Ck Can 18 049707     NCARFQEVRDTED---GNLEYCDSYFVVKQTTPSNYEHEKSCYEELKSEVTADHDFFVFN  113
               | IBV Ck EG CU 1 2014      NCARFQEVRGTED---GNFEYLDSYFVVKQTTPSNYEHEKNCYYDLKSEVTADHDFFVFN  113
               | IBV ITA 90254 2005       NCARFQEVRDTED---GNLEYCDSFFVVKQTTPSNFEHERNCYEDLKSEVTADHDFFVFN  113
                                          :*.****:    :         :*:* ****** *: **   * :**:: .* **** *:
```

nsp12 loop 67-72

```
Cegacovirus    | Beluga Whale CoV SW1     -GMYSICRQRLTKYTMMDLCYAFRHFDPNDCDVLKEILVVKGCCEWDY------------  150
Brangacovirus  | Canada Goose CoV         KSTYNIVRQRLTKYSMMDFCYALRHFDPKDCEVLKEILVDFECIPGYYSHMFGPDLSPGS  165
               | Duck CoV DK/GD/27/2014   KTIYNVSRQRLTKYTMMDLCYALRHFDPKDCEVLKEILVLNDCIEDWYWNRSGENQTPWD  176
        Anatis | Avian CoV Aus            KTIYNVSRQRLTKYTMMDLCYALRHFDPKDCEVLKEILVLNDCIEDWYWMQGGQGQAPGN  180
               | Avian CoV Grey Teal      KTIYNVSRQRLTKYTMMDLCYALRHFDPKDCEVLKEILVLNDCIEDWYWMQGGQGQAPGN  180
               | IBV Ind-TN92-03          KNIFNISRQRLTKYTKMDFCYALRHFDPKDCEVLKEILVTYGCIEDYH----------PK  163
               | IBV ck CH LLN 090312     KNIYNISRQRLTKYTMMDFCYALRHFDPKDCEVLKEILVTYGCIEDYH----------PK  163
Igacovirus     | IBV SNU 11045            KNIYNISRQRLTKYTMMDFCYALRHFDPKDCEVLKEILVTYGCIEDYH----------PK  163
               | IBV ck CH LZJ            KNIYNISRQRLTKYTMMDFCYALRHFDPKDCEVLKEILVTYGCIEDYH----------PK  163
   Galli/Pulli | Avian IBV                KNIYNISRQRLTKYTMMDFCYALRHFDPKDCEVLKEILVTYGCIEDYH----------PK  163
               | IBV Mass                 KNIYNISRQRLTKYTMMDFCYALRHFDPKDCEVLKEILVTYGCIEDYH----------PK  163
               | Turkey CoV               KNIYNISRQRLTKYTMMDFCYALRHFDPKDCEVLKEILVTYGCIEDYH----------PK  163
               | IBV Ck Can 18 049707     KNIYNISRQRLTKYTMMDFCYALRHFDPKDCEVLKEILVTYGCIEDYH----------PK  163
               | IBV Ck EG CU 1 2014      KNIYNISRQRLTKYTMMDFCYALRHFDPKDCEVLKEILVTYGCIEDYH----------PK  163
               | IBV ITA 90254 2005       KNIYNISRQRLTKYTMMDFCYALRHFDPKDCEVLKEILVTYGCIEDYH----------PK  163
                                             :.: *******: **:***:*****:**:*******   *    :
```

nsp12 loop 113-115

nsp12 loop 156-169

```
Cegacovirus    | Beluga Whale CoV SW1     --FDQPNWYDPVENPDWFSLISRLGPIFQRALIKVAEFCDLMVEKGYIGVVTLDNQDLNG  208
Brangacovirus  | Canada Goose CoV         WFDHNPLWYDPIENPSYHDIMANLGCVVNRAVLNCVKFCDLMVDKGYIGVITLDNQDMNG  225
               | Duck CoV DK/GD/27/2014   WFDHNLTWYDPIENPSYYAILANLGHIVRHALLNAVKFGDLMVEKGYVGVVTLDNQDLNG  236
        Anatis | Avian CoV Aus            WFDHNPTWYDPIENPSYYAILANLGHIVRRALLNAVKFGDLMVEKGYVGVVTLDNQDLNG  240
               | Avian CoV Grey Teal      WFDHNPTWYDPIENPSYYAILANLGHIVRRALLNAVKFGDLMVEKGYVGVVTLDNQDLNG  240
               | IBV Ind-TN92-03          WFEENKDWYDPIENPKYYAMLAKMGPIVRRALLNAIEFSNLMVEKGYVGVITLDNQDLNG  223
               | IBV ck CH LLN 090312     WFEENKDWYDPVENPKYYAMWANMGPIVRRALLNAVEFGNLMVDKGYVGVVTLDNQDLNG  223
Igacovirus     | IBV SNU 11045            WFEENKDWYDPIENPKYYAMLAKMGPIVRRALLNAVEFGNLMVEKGYVGVVTLDNQDLNG  223
               | IBV ck CH LZJ            WFEENKDWYDPIENPKYYAMLAKMGPIVRRALLNAVEFGNLMVEKGYVGVVTLDNQDLNG  223
   Galli/Pulli | Avian IBV                WFEENKDWYDPIENSKYYVMLAKMGPIVRRALLNAIEFGNLMVEKGYVGVITLDNQDLNG  223
               | IBV Mass                 WFEENKDWYDPIENPKYYAMLAKMGPIVRRALLNAIEFGNLMVEKGYVGVITLDNQDLNG  223
               | Turkey CoV               WFEENKDWYDPIENPKYYAMLAKVGPIVRRALLNAIEFGNLMVEKGYVGVVTLDNQDLNG  223
               | IBV Ck Can 18 049707     WFEENKDWYDPIENPKYYAMLAKMGPIVRRALLNAIEFGNLMVEKGYVGVITLDNQDLNG  223
               | IBV Ck EG CU 1 2014      WFEENKDWYDPIENPKYYAMLAKMGPIVRRALLNAIEFGTLMVEKGYVGVVTLDNQDLNG  223
               | IBV ITA 90254 2005       WFEENKDWYDPIENPKYYAMLAKMGPIVRRALLNAIEFGNFMVEKGYVGVVTLDNQDLNG  223
                                             .:  ****:** .:. : :.:* :..:*::   :*  :**:***:**:******:**
```

```
Cegacovirus    | Beluga Whale CoV SW1     NFYDFGDFKKVLPGCGVPVTTSYYSYMMPCLTACDALASERFFEFKATSGYKQYDLTKYD  268
Brangacovirus  | Canada Goose CoV         KFYDFGDFQKTMPGSGVPVVDSYYSYLMPLLTMTDALACERYYNFDKIKGFKTYDLFKYD  285
               | Duck CoV DK/GD/27/2014   KFYDFGDFQKTMPGAGVPIFDSYYSYIMPIITMTDALATERFYDFDRHKGFKSYDLLKYD  296
        Anatis | Avian CoV Aus            KFYDFGDFQKTTPGAGVPIFDSYYSYLMPIITMTDALATERFYEFDRHKGFKSYDLLKYD  300
               | Avian CoV Grey Teal      KFYDFGDFQKTTPGAGVPIFDSYYSYLMPIITMTDALATERFYEFDRHKGFKSYDLLKYD  300
               | IBV Ind-TN92-03          KFYDFGDFQKTAPGAGVPVFDTYYSYKMPIIAKTDAFAPERYFEYDVHKGYKSYDLLKYD  283
               | IBV ck CH LLN 090312     KFYDFGDFQKTAPGAGVPVFDTYYSYMMPIIAMTDALAPERYFEHDVHKGYKSYDLLKYD  283
Igacovirus     | IBV SNU 11045            KFYDFGDFQKTAPGAGVPVFDTYYSYMMPIIAMTDALAPERYFEYDVHKGYKSYDLLKYD  283
               | IBV ck CH LZJ            KFYDFGDFQKTAPGAGVPVFDTYYSYMMPIIAMTDALAPERYFEYDVHKGYKSYDLLKYD  283
   Galli/Pulli | Avian IBV                KFYDFGDFQKTAPGAGVPVFDTYYSYMMPIIAMTDALAPERYFEYDVHKGYKSYDLLKYD  283
               | IBV Mass                 KFYDFGDFQKTAPGAGVPVFDTYYSYMMPIIAMTDALAPERYFEYDVHKGYKSYDLLKYD  283
               | Turkey CoV               KFYDFGDFQKTALGAGIPVFDTYYSYMMPIIAMTDALAPERYFEYDVHKGYKSYDLLKYD  283
               | IBV Ck Can 18 049707     KFYDFGDFQKTALGAGVPVFDTYYSYMMPIIAMTDALAPERYFEYDVHKGYKSYDLLKYD  283
               | IBV Ck EG CU 1 2014      KFYDFGDFQKTAPGAGVPVFDTYYSYMMPIIAMTNALAPERYFEYDVHKGYKSYDLLKYD  283
               | IBV ITA 90254 2005       KFYDFGDFQKTAPGAGVPVFDTYYSYMMPIIAMTDALAPERYFEYDVHKGYKSYDLLKYD  283
                                          :*******:*.  *.*:*:  :**** ** ::  :*:* **:::..  .*:* *** ***
```

nsp12 loop 264-278

```
Cegacovirus   | Beluga Whale CoV SW1     FTEEKLQLFMKYFKYWDRTYHPNCVECIDDRCLIHCANFNILFATLFPQTAFGCLCKRVY   328
Brangacovirus | Canada Goose CoV         YTEEKRQLFNKYFKYWDQEYHPNCCDCVDDRCVIHCANFNVLFSTLIPQTSFGNLCRKVY   345
                Duck CoV DK/GD/27/2014   YTEEKLELFSKYFKYWDQEYHPNCSECADDRCLIHCANFNILFSTLIPQTSFGNLCRKVY   356
        Anatis  Avian CoV Aus            YTEEKLELFSKYFKYWDQEYHPNCSECADDRCLIHCANFNILFSTLIPQTSFGNLCRKVY   360
                Avian CoV Grey Teal      YTEEKLELFSKYFKYWDQEYHPNCSECADDRCLIHCANFNILFSTLIPQTSFGNLCRKVY   360
                IBV Ind-TN92-03          YTEEKQELFQKYFKYWDQEYHPNCRDCSDDRCLIHCAYFTILFSRLVPQTSFGNLCRKVF   343
                IBV ck CH LLN 090312     YTEEKQELFQKYFKYWDQEYHPNCRDCVDDRCLIHCANFNVLFSTLIPQTSFGNLCRKVF   343
Igacovirus      IBV SNU 11045            YTEEKQALFQKYFKYWDQEYHPNCRDCVDDRCLIHCANFNVLFSTLIPQTSFGNLCRKVF   343
                IBV ck CH LZJ            YTEEKQALFQKYFKYWDQEYHPNCRDCVDDRCLIHCANFNVLFSTLIPQTSFGNLCRKVF   343
   Galli/Pulli  Avian IBV                YTEEKQELFQKYFKYWDQEYHPNCRDCSDDRCLIHCANFNILFSTLIPQTSFGNLCRKVF   343
                IBV Mass                 YTEEKQDLFQKYFKYWDQEYHPNCRDCSDDRCLIHCANFNILFSTLVPQTSFGNLCRKVF   343
                Turkey CoV               YTEEKQEMFQKYFKYWDQEYHPNCRDCSDDRCLIHCANFNILFSTLIPQTSFGNLCRKVF   343
                IBV Ck Can 18 049707     YTEEKQEMFQKYFKYWDQEYHPNCRDCSDDRCLIHCANFNILFSTLIPQTSFGNLCRKVF   343
                IBV Ck EG CU 1 2014      YTEEKQELFQKYFKYWDQEYHPNCRDCSDDRCLIHCANFNILFSTLIPQTSFGNLCRKVF   343
                IBV ITA 90254 2005       YTEEKQELFQKYFKYWDQEYHPNCRDCSDDRCLIHCANFNILFSTLIPQTSFGNLCRKVF   343
                                         :****  :* *******: ***** :* ****:**** *.:**: *.***:** **::*:

Cegacovirus   | Beluga Whale CoV SW1     IDGVPFISTTGYHSKELGVLLNKDNSMSFSKMSIGELMRFAADPSLLVSASDAFVDLRTS   388
Brangacovirus | Canada Goose CoV         VDGVPFIATTGYHSKELGVIMNQDNTLSFSKMSLTQLMKFVGDPALLVGTSNNLVDLRTS   405
                Duck CoV DK/GD/27/2014   VDGVPFIATCGYHSKELGVIMNQDNTLSFSKMGLGQLMKFVGDPALLVGTSNNLVDLRTS   416
        Anatis  Avian CoV Aus            VDGVPFIATCGYHSKELGVIMNQDNTLSFSKMGLGQLMKFVGDPALLVGTSNNLVDLRTS   420
                Avian CoV Grey Teal      VDGVPFIATCGYHSKELGVIMNQDNTLSFSKMGLGQLMKFVGDPALLVGTSNNLVDLRTS   420
                IBV Ind-TN92-03          VDGVPFIATCGYHSKELGVIKNQDNTMSFSKMGLCQLMQFVGDPALFVGTSNNFVDLRTS   403
                IBV ck CH LLN 090312     VDGVPFIATCGYHSKELGVIMNQDNTMSFSKMGLSQLMKFVGDPALLVGTSNNLVDLRTS   403
Igacovirus      IBV SNU 11045            VDGVPFIATCGYHSKELGVIMNQDNTMSFSKMGLSQLMKFVGDPALLVGTSNNLVDFRTS   403
                IBV ck CH LZJ            VDGVPFIATCGYHSKELGVIMSQDNTMSFSKMGLSQLMKFVGDPALLVGTSNNLVDLRTS   403
   Galli/Pulli  Avian IBV                VDGVPFIATCGYHSKELGVIMNQDNTMSFSKMGLSQLMQFVGDPALLVGTSNNLVDLRTS   403
                IBV Mass                 VDGVPFIATCGYHSKELGVIMNQDNTMSFSKMGLSQLMQFVGDPALLVGTSNKLVDLRTS   403
                Turkey CoV               VDGVPFIATCGYHSKELGVIMNQDNTMSFSKLGLSQLMQFVGDPALLVGTSNNLVDLRTS   403
                IBV Ck Can 18 049707     VDGVPFIATCGYHSKELGVIMNQDNTMSFSKMGLSQLMQFVGDPALLVGTSNNLVDLRTS   403
                IBV Ck EG CU 1 2014      VDGVPFIATCGYHSKELGVIMNQDNTMSFSKMGLSQLMQFVGDPALLVGTSNNLVDLRTS   403
                IBV ITA 90254 2005       VDGVPFIATCGYHSKELGVIMNQDNTMSFSKMGLSQLMQFVGDPALLVGTSNNLVDLRTS   403
                                         :******:* *********: .:**::****:.: :**:*..**:*:*.:*: :**:***

Cegacovirus   | Beluga Whale CoV SW1     CFSLSALSTGLTYQTVKPGHFNEDFYNFAEKKGFFKEGSSIPLKHFFYIQDGNAAIADFD   448
Brangacovirus | Canada Goose CoV         CFSLCALVTGITHQTVKPGHFNRDFYDFAEKSGMFKEGSSIPLKHFFFPQTGSAAIEDYD   465
                Duck CoV DK/GD/27/2014   CFSICALASGITHQTVKPGHFNKDFYDFAEKAGMFKEGSSIPLKHFFFPQTGSAAIDDYD   476
        Anatis  Avian CoV Aus            CFSICALASGITHQTVKPGHFNKDFYDFAEKAGMFKEGSSIPLKHFFFPQTGSAAIDDYD   480
                Avian CoV Grey Teal      CFSICALASGITHQTVKPGHFNKDFYDFAEKAGMFKEGSSIPLKHFFFPQTGSAAIDDYD   480
                IBV Ind-TN92-03          CFSVCALSSGITHQTVKPGHFNKDFYDSAEKAGMFKEGSSLPLKHFFYPQTGNAAIYDYE   463
                IBV ck CH LLN 090312     CFSVCALASGITHQTVKPGHFNKDFYDFAEKAGMFKEGSSIPLKHFFYPQTGNAAINDYD   463
Igacovirus      IBV SNU 11045            CFSVCALASGITHQTVKPGHFNKDFYDFAEKAGMFKEGSSIPLKHFFYPQTGNAAINDYD   463
                IBV ck CH LZJ            CFSVCALASGITHQTVKPGHFNKDFYDFAEKAGMFKEGSSIPLKHFFYPQTGNAAINDYD   463
   Galli/Pulli  Avian IBV                CFSVCALTSGITHQTVKPGHFNKDFYDFAEKAGMFKEGSSIPLKHFFYPQTGNAAINDYD   463
                IBV Mass                 CFSVCALASGITHQTVKPGHFNKDFYDFAEKAGMFKEGSSIPLKHFFYPQTGNAAINDYD   463
                Turkey CoV               CFSVCALASGITHQTVKPGHFNKDFYDFAEKAGMFKEGSSIPLKHFFYPQTGSAAINDYD   463
                IBV Ck Can 18 049707     CFSVCALASGITHQTVKPGHFNKDFYDFAEKAGMFKEGSSIPLKHFFYPQTGNAAINDYD   463
                IBV Ck EG CU 1 2014      CFSVCALASGITHQTVKPGHFNKDFYDFAEKAGMFKEGSSIPLKHFFYPQTGNAAINDYD   463
                IBV ITA 90254 2005       CFSVCALASGVTHQTVKPGHFNKDFYDFAEKAGMFKEGSSIPLKHFFYPQTGNAAINDYD   463
                                         ***:.** :*:*:*********.***: *** *:******:******: * *.*** *::

Cegacovirus   | Beluga Whale CoV SW1     YYRFNKPTMVDIQQFLFCFEVTDKYFEQYDGGCIPANQVVVTNLDKSAGYPFNKFGKARL   508
Brangacovirus | Canada Goose CoV         YYRYNRPTMFDIRQLLFCLEVTAKYFDCYEGGCIPASQVIVTNLDKSAGYPFNKFGKARL   525
                Duck CoV DK/GD/27/2014   YYRYNRHTMFDIRQLLFCLEVTSKYFECYEGGCISASQVVVTNLDKSAGYPFNKFGKARL   536
        Anatis  Avian CoV Aus            YYRYNRPTMFDIRQLLFCLEVTSKYFECYEGGCISASQVVVTNLDKSAGYPFNKFGKARL   540
                Avian CoV Grey Teal      YYRYNRPTMFDIRQLLFCLEVTSKYFECYEGGCISASQVVVTNLDKSAGYPFNKFGKARL   540
                IBV Ind-TN92-03          YYRYIRPTMFDICQILFCFEETSKYFECYEGGCIPAGQVLVNNFDKSPGYPFNKFGKARL   523
                IBV ck CH LLN 090312     YYRYNRPTMFDIRQLLFCLEVTSKYFECYEGGCIPASQVVVTNLDKSAGFPFNKFGKARL   523
Igacovirus      IBV SNU 11045            YYRYNRPTMFDIRQLLFCLEVTSKYFECYEGGCIPASQVVVTNLDKSAGFPFNKFGKARL   523
                IBV ck CH LZJ            YYRYNRPTMFDIRQLLFCLEVTSKYFECYEGGCIPASQVVVTNLDKSAGFPFNKFGKARL   523
   Galli/Pulli  Avian IBV                YYRYNRPTMFDICQLLFCLEVTSKYFECYEGGCIPASQVVVNNLDKSAGYPFNKFGKARL   523
                IBV Mass                 YYRYNRPTMFDIRQLLFCLEVTSKYFECYEGGCIPASQVVVNNLDKSAGYPFNKFGKARL   523
                Turkey CoV               YYRYNRPTMFDIRQLLFCLEVTSKYFDCYEGGCIPASQVVVNNLDKSAGYPFNKFGKARL   523
                IBV Ck Can 18 049707     YYRYNRPTMFDIRQLLFCLEVTSKYFECYEGGCIPASQVVVNNLDKSAGYPFNKFGKARL   523
                IBV Ck EG CU 1 2014      YYRYNRPTMFDVRQLLFCLEVTSKYFECYEGGCIPASQVVVNNLDKSAGYPFNKFGKARL   523
                IBV ITA 90254 2005       YYRYNRPTMFDIRQLLFCLEVTSKYFDCYEGGCIPASQVVVNNLDKSAGYPFNKFGKARL   523
                                         ***: : **.*: *:***:* * ***: *:**** *.**:*.*:*** *:******:***

Cegacovirus   | Beluga Whale CoV SW1     YYESLSYAEQDQLFELTKRNILPTITQINMKYAISAKSRARTVAGVSILSSMTNRQFHQK   568
Brangacovirus | Canada Goose CoV         YYE-MSYEEQDALFASTKRNVMPTITQMNLKYAISAKNRARTVAGVSILSTMTNRQYHQK   584
                Duck CoV DK/GD/27/2014   YYE-MSFEEQDQLFESTKKNVLPTITQMNLKYAISAKNRARTVAGVSILSTMTNRQYHQK   595
        Anatis  Avian CoV Aus            YYE-MSFEEQDQLFESTKKNVLPTITQMNLKYAISAKNRARTVAGVSILSTMTNRQYHQK   599
                Avian CoV Grey Teal      YYE-MSFEEQDQLFESTKKNVLPTITQMNLKYAISAKNRARTVAGVSILSTMTNRQYHQK   599
                IBV Ind-TN92-03          YYE-KSLEEQDQLFESTKKNVLPTISQKNLKYAISAKNRARTVAGVSILSTKTNRQFHQK   582
                IBV ck CH LLN 090312     YYE-MSLEEQDQLFESTKKNVLPTITQMNLKYAISAKNRARTVAGVSILSTMTNRQFHQK   582
Igacovirus      IBV SNU 11045            YYE-MSLEEQDQLFESTKKNVLPTITQMNLKYAISAKNRARTVAGVSILSTMTNRQFHQK   582
                IBV ck CH LZJ            YYE-MSLEEQDQLFESTKKNVLPTITQMNLKYAISAKNRARTVAGVSILSTMTNRQFHQK   582
   Galli/Pulli  Avian IBV                YYE-MSLEEQDQLFEITKKNVLPTITQMNLKYAISAKNRARTVAGVSILSTMTNRQFHQK   582
                IBV Mass                 YYE-MSLEEQDQLFESTKKNVLPTITQMNLKYAISAKNRARTVAGVSILSTMTNRQFHQK   582
                Turkey CoV               YYE-MSLEEQDQLFESTKKNVLPTITQMNLKYAISAKNRARTVAGVSILSTMTNRQFHQK   582
                IBV Ck Can 18 049707     YYE-MSLEEQDQLFESTKKNVLPTITQMNLKYAISAKNRARTVAGVSILSTMTNRQFHQK   582
                IBV Ck EG CU 1 2014      YYE-MSLEEQDQLFESTKKNVLPTITQMNLKYAISAKNRARTVAGVSILSTMTNRQFHQK   582
                IBV ITA 90254 2005       YYE-MSLDEQDQLFESTKKNVLPTITQMNLKYAISAKNRARTVAGVSILSTMTNRQFHQK   582
                                         ***  *  *** **  **:*::***:* *:*******.************: ****:***
```

```
Cegacovirus   |               Beluga Whale CoV SW1      CLKSIVNTRNATVVIGTTKFYGGWDNMLRNLMRGVEDPVLMGWDYPKCDRAVPSLLRLSA   628
Brangacovirus |               Canada Goose CoV          VLKSIVNTRNAAVVIGTSKFYGGWDNMLRNLIDGVDNPMLMGWDYPKCDRAMPNILRISA   644
              |               Duck CoV DK/GD/27/2014    VLKSIVNTRNAPVVIGTTKFYGGWDNMLRNLVKGVDDPILMGWDYPKCDRAMPNLLRIAA   655
              | Anatis        Avian CoV Aus             VLKSIVNTRNAPVVIGTTKFYGGWDNMLRNLIKGVDDPILMGWDYPKCDRAMPNLLRIAA   659
              |               Avian CoV Grey Teal       VLKSIVNTRNAPVVIGTTKFYGGWDNMLRNLIKGVDDPILMGWDYPKCDRAMPNLLRIAA   659
              |               IBV Ind-TN92-03           ILKSIVNTRNAPVVIGTTKFYGGWDNMLRNLIQGVEDPILMGWDYPKCDRAMPNLLRIAA   642
              |               IBV ck CH LLN 090312      VLKSIVNTRNAPVVIGTTKFYGGWDNMLRNLVQGVEDPMLMGWDYPKCDRAMPNLLRIAA   642
Igacovirus    |               IBV SNU 11045             VLKSIVNTRNAPVVIGTTKFYGGWDNMLRNLVQGVEDPMLMGWDYPKCDRAMPNLLRIAA   642
              |               IBV ck CH LZJ             VLKSIVNTRNAPVVIGTTKFYGGWDNMLRNLVQGVEDPMLMGWDYPKCDRAMPNLLRIAA   642
              | Galli/Pulli   Avian IBV                 ILKSIVNTRNASVVIGTTKFYGGWDNMLRNLIQGVEDPILMGWDYPKCDRAMPNLLRIAA   642
              |               IBV Mass                  ILKSIVNTRNAPVVIGTTKFYGGWDNMLRNLIQGVEDPILMGWDYPKCDRAMPNLLRIAA   642
              |               Turkey CoV                ILKSIVNTRNAPVVIGTTKFYGGWDNMLRNLIQGVEDPILMGWDYPKCDRAMPNLLRIAA   642
              |               IBV Ck Can 18 049707      ILKSIVNTRNAPVVIGTTKFYGGWDNMLRNLIQGVEDPILMGWDYPKCDRAMPNLLRIAA   642
              |               IBV Ck EG CU 1 2014       ILKSIVNTRNASVVIGTTKFYGGWDNMLRNLIQGVEDPILMGWDYPKCDRAMPNLLRIAA   642
              |               IBV ITA 90254 2005        ILKSIVNTRNAPVVIGTTKFYGGWDNMLRNLIQGVENPILMGWDYPKCDRAMPNLLRIAA   642
                                                         ********** *****:*************: **::*:************:*.:**::*

Cegacovirus   |               Beluga Whale CoV SW1      SLILARRHKGCCDWNERIYRLANEAAQVLSEVALSNGGLYVKPGGTSSGDATTAYANSAF   688
Brangacovirus |               Canada Goose CoV          SLVLARKHTTCCTWSERIYRLFNECAQVLSESVLATGGIYVKPGGTSSGDATTAYANSVF   704
              |               Duck CoV DK/GD/27/2014    SLVLARKHTNCCTWSDRVYRLYNECAQVLSESVLATGGIYVKPGGTSSGDATTAYANSVF   715
              | Anatis        Avian CoV Aus             SLVLARKHTNCCTWSDRVYRLYNECAQVLSESVLATGGIYVKPGGTSSGDATTAYANSVF   719
              |               Avian CoV Grey Teal       SLVLARKHTNCCTWSDRVYRLYNECAQVLSESVLATGGIYVKPGGTSSGDATTAYANSVF   719
              |               IBV Ind-TN92-03           SFVLARKHTICCTWSERVYWLYIECAQVLSETVLATGGIYVKPGGTCSGDATTAYANSVF   702
              |               IBV ck CH LLN 090312      SLVLARKHTNCCTWSERIYRLYNECAQVLSETVLATGGIYVKPGGTSSGDATTAYANSVF   702
Igacovirus    |               IBV SNU 11045             SLVLARKHTNCCTWSERIYRLYNECAQVLSETVLATGGIYVKPGGTSSGDATTAYANSVF   702
              |               IBV ck CH LZJ             SLVLARKHTNCCTWSERIYRLYNECAQVLSETVLATGGIYVKPGGTSSGDATTAYANSVF   702
              | Galli/Pulli   Avian IBV                 SLVLARKHTNCCSWSERIYRLYNECAQVLSETVLATGGIYVKPGGTSSGDATTAYANSVF   702
              |               IBV Mass                  SLVLARKHTNCCTWSERVYRLYNECAQVLSETVLATGGIYVKPGGTSSGDATTAYANSVF   702
              |               Turkey CoV                SLVLARKHTNCCTWSERIYRLYNECAQVLSETVLATGGIYVKPGGTSSGDATTAYANSVF   702
              |               IBV Ck Can 18 049707      SLVLARKHTNCCTWSERIYRLYNECAQVLSETVLATGGIYVKPGGTSSGDATTAYANSVF   702
              |               IBV Ck EG CU 1 2014       SLVLARKHTNCCTWPERIYRLYNECAQVLSETVLATGGIYVKPGGTSSGDATTAYANSVF   702
              |               IBV ITA 90254 2005        SLVLARKHTNCCTWSERIYRLYNECAQVLSETVLATGGIYVKPGGTSSGDATTAYANSVF   702
                                                        *::***:*. ** * :*:* *  *.****** .*:.**:*******.***********.*

Cegacovirus   |               Beluga Whale CoV SW1      NLFQATAANVAQLLATPTSRIYAEEVRALQHELYTQVYRRDKPDMDFVYTFYAYLNKHFS   748
Brangacovirus |               Canada Goose CoV          NIIQATSANVARLISTCTPDIVYDDVRSMQYELYQQVYRRVNPDPAFVDKFFAYLNKHFS   764
              |               Duck CoV DK/GD/27/2014    NIIQATSANVARLVSTITRDIVYDDVRAMQHELYQQVYRRVNPDAAFVEKFYSYLNKHFS   775
              | Anatis        Avian CoV Aus             NIIQATSANVARLVSTITRDIVYDDVRAMQHELYQQVYRRVNPDAAFVEKFYSYLNKHFS   779
              |               Avian CoV Grey Teal       NIIQATSANVARLVSTITRDIVYDDVRAMQHELYQQVYRRVNPDASFVEKFYSYLNKHFS   779
              |               IBV Ind-TN92-03           NIIQATSAIVARLLSVITRDIVYEDIMSLQYELYQQVYWRVNFDPAFVEKFYSYFCKNFS   762
              |               IBV ck CH LLN 090312      NIIQATSANVARLLSAITRDIVYDDIKSLQYELYQQVYRRVNFDPAFVEKFYFYMCKNFS   762
Igacovirus    |               IBV SNU 11045             NIIQATSANVARLLSVITRDIVYDDIKSLQYELYQQVYRRVNFDPSFVEKFYSYMCKNFS   762
              |               IBV ck CH LZJ             NIIQATSANVARLLSVITRDIVYDDIKSLQYELYQQVYRRVNFDPSFVEKFYFYMCKNFS   762
              | Galli/Pulli   Avian IBV                 NIIQATSANVARLLSVITRDIVYDNIKSLQYELYQQVYRRVNFDPAFVEKFYSYLCKNFS   762
              |               IBV Mass                  NIIQATSANVARLLSVITRDIVYDDIKSLQYELYQQVYRRVNFDPAFVEKFYSYLCKNFS   762
              |               Turkey CoV                NIIQATSANVARLLSVITRDIVYDDIKSLQYELYQQVYRRVNFDPAFVEKFYSYLCKNFS   762
              |               IBV Ck Can 18 049707      NIIQATSANVARLLSVITRDIVYDDIKSLQYELYQQVYRRVNFDPAFVEKFYSYLCKNFS   762
              |               IBV Ck EG CU 1 2014       NIIQATSANVARLLSVITRDIVYDDIKDLQYELYQQVYRRVNFDSAFVEKFYSYLCKNFS   762
              |               IBV ITA 90254 2005        NIIQATSANVARLLSVITRDIVYDDIKSLQYELYQQVYRRVNLDPAFVEKFYSYLCKNFS   762
                                                        *::***:* **:*::. *  *  :::  :*:*** *** * : *  ** .*: *: *:**

Cegacovirus   |               Beluga Whale CoV SW1      LMILSDDGVVCYNKSYAEAGMVASIASFREVLFYQNNVFMADSKCWTEEDVKIGPHEFCS   808
Brangacovirus |               Canada Goose CoV          LMILSDDGVVCYNDTYAKQGLVADISAFREVLYYQNNVFMSDAKCWTEPDIIKGPHEFCS   824
              |               Duck CoV DK/GD/27/2014    LMILSGDGVVCYNNTYAKQGLVADISAFREVLYYQNNVYMADSKCWVEPDLEKGPHEFCS   835
              | Anatis        Avian CoV Aus             LMILSDDGVVCYNNTYAKQGLVADISAFREVLYYQNNVYMADSKCWVEPDLEKGPHEFCS   839
              |               Avian CoV Grey Teal       LMILSDDGVVCYNNTYAKQGLVADISAFREVLYYQNNVYMADSKCWVEPDLEKGPHEFCS   839
              |               IBV Ind-TN92-03           LMILSDDGVVCYNNTLAKQGLVADISGFREVLYYQNNVFMADSKCWVEPDLEKGPHEFCS   822
              |               IBV ck CH LLN 090312      LMILSDDGVVCYNNTLARQGLVADISGFREILYYQNNVYMADSKCWVEPDLEKGPHEFCS   822
Igacovirus    |               IBV SNU 11045             LMILSDDGVVCYNNTLAKQGLVADISGFREILYYQNNVYMADSKCWVEPDLENGPHEFCS   822
              |               IBV ck CH LZJ             LMILSDDGVVCYNNTLAKQGLVADISGFREILYYQNNVYMADAKCWVEPDLKKGPHEFCS   822
              | Galli/Pulli   Avian IBV                 LMILSDDGVVCYNNTLAKQGLVADISGFREVLYYQNNVFMADSKCWVEPDLEKGPHEFCS   822
              |               IBV Mass                  LMILSDDGVVCYNNTLAKQGLVADISGFREVLYYQNNVFMADSKCWVEPDLEKGPHEFCS   822
              |               Turkey CoV                LMILSDDGVVCYNNTLAKQGLVADISGFREILYYQNNVYMADSKCWVEPDLEKGPHEFCS   822
              |               IBV Ck Can 18 049707      LMILSDDGVVCYNNTLAKQGLVADISGFREVLYYQNNVFMSDSKCWVEPDLEKGPHEFCS   822
              |               IBV Ck EG CU 1 2014       LMVLSDDGVVCYNNTLAKQGLVADISGFREILYYQNNVYMADSKCWVEPDLEKGPHEFCS   822
              |               IBV ITA 90254 2005        LMILSDDGVVCYNNTLAKQGLVADISGFREILYYQNNVYMADSKCWVEPDLEKGPHEFCS   822
                                                        **:**.*******.: *. *:**.*:.***:*:*****:*:*:***.* *:  *******

Cegacovirus   |               Beluga Whale CoV SW1      QHSMLVEIDGEMRYLPYPDPSRILGACVFVDDVEKTEPVVVMERYVALAIDAYPLIYHEN   868
Brangacovirus |               Canada Goose CoV          QHTMLVEVDGEYKYLPYPDPSRILGACVFVDDIEKTEPVAVMERYVALAIDAYPLVFHEN   884
              |               Duck CoV DK/GD/27/2014    QHTMLVDVDGEPKYLPYPDPSRILGACVFVDDVDKTEPVAVMERYVALAIDAYPLVYHEN   895
              | Anatis        Avian CoV Aus             QHTMLVDVDGEPKYLPYPDPSRILGACVFVDDVDKTEPVAVMERYVALAIDAYPLVYHEN   899
              |               Avian CoV Grey Teal       QHTMLVDVDGEPRYLPYPDPSRILGACVFVDDVDKTEPVAVMERYVALAIDAYPLVYHEN   899
              |               IBV Ind-TN92-03           QHTMLVEVDGEPRYLPYPDPSRILCACVFVGDLDKTESVAVMERYIALAIDAYPLVHHEN   882
              |               IBV ck CH LLN 090312      QHTMLVEVDGEPRYLPYPDPSRILGSCVFVDEVDKTEPVAVMERYIALAIDAYPLVHHEN   882
Igacovirus    |               IBV SNU 11045             QHTMLVEVDGEPRYLPYPDPSRILGACVFVDEVDKTEPVAVMERYIALAIDAYPLVHHEN   882
              |               IBV ck CH LZJ             QHTMLVEVDGEPRYLPYPDPSRILGACVFVDEVDKTEPVAVMERYIALAIDAYPLVHHEN   882
              | Galli/Pulli   Avian IBV                 QHTMLVEVDGEPKYLPYPDPSRILGACVFVDDVDKTEPVAVMERYIALAIDAYPLVHHEN   882
              |               IBV Mass                  QHTMLVEVDGEPRYLPYPDPSRILCACVFVDDLDKTESVAVMERYIALAIDAYPLVHHEN   882
              |               Turkey CoV                QHTMLVEVDGEPKYLPYPDPSRILGACVFVDDVDKTEPVAVMERYIALAIDAYPLVHHEN   882
              |               IBV Ck Can 18 049707      QHTMLVEVDGEPKYLPYPDPSRILGACVFVDDVDKTEPVAVMERYIALAIDAYPLVHHEN   882
              |               IBV Ck EG CU 1 2014       QHTMLVEVDGEPKYLPYPNPSRILGACVFVDEVDKTEPVAVMERYIALAIDAYPLVHHEN   882
              |               IBV ITA 90254 2005        QHTMLVEVDGEPKYLPYPDPSRILGACVFVDEVDKTEPVAVMERYIALAIDAYPLVHHEN   882
                                                        **:***::*** :*****:***** :****.:::*** *.*****:*********:.***
```

```
  Cegacovirus  | Beluga Whale CoV SW1     EEYGKVFYLLLSYIQTLYQRLSNDMLMDYSFVMNIDTSSKFWEEDFYRQMYGSSPTLQ     926
Brangacovirus  | Canada Goose CoV         EEYRRVFYVLLAYIRKLYQELSKNMLTDYSFVLDIDKGSKFWEEEFYSNMYRTPTTLQ     942
Igacovirus
       Anatis      | Duck CoV DK/GD/27/2014   EEYKKVFYVLLSYIRKLYQELSKNMLTDYSFVLDIDKGSKFWEEEFYSNMYRAPTTLQ     953
                   | Avian CoV Aus            EEYKKVFYVLLSYIRKLYQELSKNMLTDYSFVLDIDKGSKFWEEEFYSNMYRAPTTLQ     957
                   | Avian CoV Grey Teal      EEYKKVFYVLLSYIRKLYQELSKNMLTDYSFVLDIDKGSKFWEEEFYSNMYRAPTTLQ     957
       Galli/Pulli | IBV Ind-TN92-03          EEYKKVFFVLLSYIRKLYQELSQNMLMDYSFVMDIDKGSKFWEQEFYENMYRAPTTLQ     940
                   | IBV ck CH LLN 090312     EEYKKVFFVLLSYIRKLYQELSQSMLIDYSFVMDIDKGSKFWEQEFYENMYRAPTTLQ     940
                   | IBV SNU 11045            EEYKKVFFVLLSYIRKLYQELSQSMLIDYSFVMDIDKGSKFWEQEFYENMYRAPTTL-     939
                   | IBV ck CH LZJ            EEYKKVFFVLLSYIRKLYQELSQSMLIDYSFVMDIDKGSKFWEQEFYENMYRAPTTLQ     940
                   | Avian IBV                EEYKKVFFVLLAYIRKLYQELSQNMLMDYSFVMDIDKGSKFWEQEFYENMYRAPTTLQ     940
                   | IBV Mass                 EEYKKVFFVLLSYIRKLYQELSQNMLMDYSFVMDIDKGSKFWEQEFYENMYRAPTTLQ     940
                   | Turkey CoV               EEYKKVFFVLLSYIRKLYQELSQNMLMDYSFVMDIDKGSKFWEQEFYENMYRAPTTLQ     940
                   | IBV Ck Can 18 049707     EEYKKVFFVLLSYIRKLYQELSQNMLMDYSFVMDIDKGSKFWEQEFYENMYRAPTTLQ     940
                   | IBV Ck EG CU 1 2014      EEYRKVFFVLLSYIRKLYQELSQSMLMDYSFVMDIDKGSKFWEQEFYENMYRAPTTLQ     940
                   | IBV ITA 90254 2005       EEYRKVFFVLLSYIRKLYQELSQSMLMDYSFVMDIDKGSKFWEQEFYENMYRAPTTLQ     940
                                            *** :**::**:**:.***.**:.** *****::**..*****::** :** :  **
```

**Figure S11 Gammacoronavirus genus nsp12 sequence alignment**: Clustal omega alignment of nsp12 proteins from viruses residing within the gammacoronavirus genus. “*” notation means residue is fully conserved, “:” means residue is strongly conserved, and “.” Denotes a weakly conserved residue.

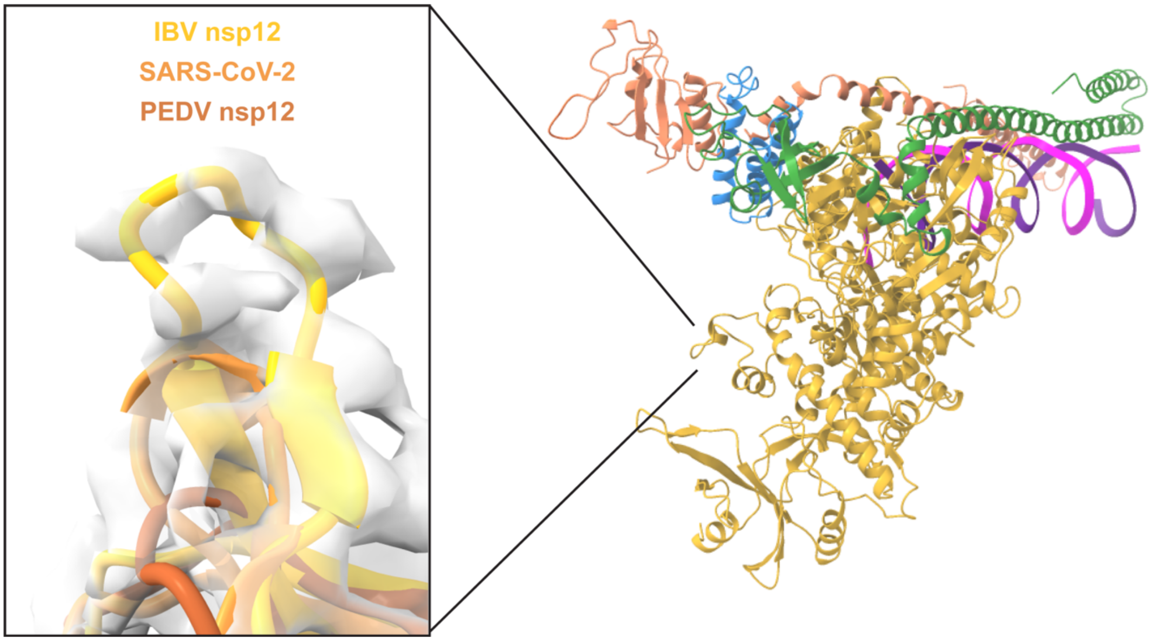


**Figure S12 Structural view of IBV nsp12 67-72 insertion:** Cartoon model of the IBV complex (right) with zoomed in view of superimposed models of IBV, PEDv (8URB) and SARS-CoV-2 (6XEZ). Focused view on IBV nsp12 67-72 insertion with electron density to support inserted region conformation.

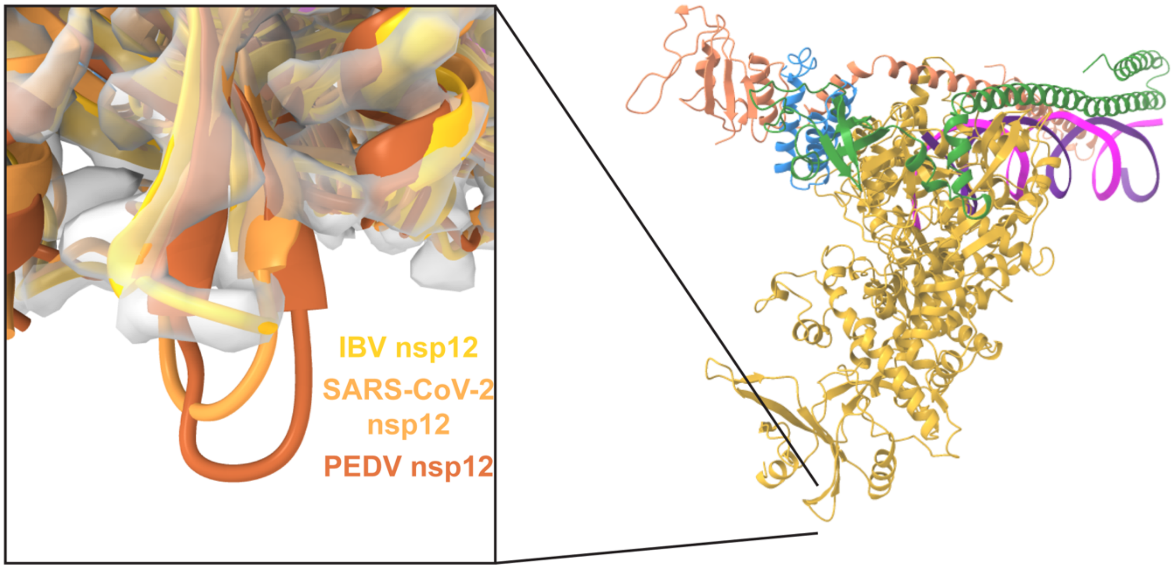


**Figure S13 Structural view of IBV nsp12 113-115 shortened loop:** Cartoon model of the IBV complex (right) with zoomed in view of superimposed models of IBV, PEDv (8URB) and SARS-CoV-2 (6XEZ). Focused view on IBV nsp12 shortened loop with electron density to support loop conformation.

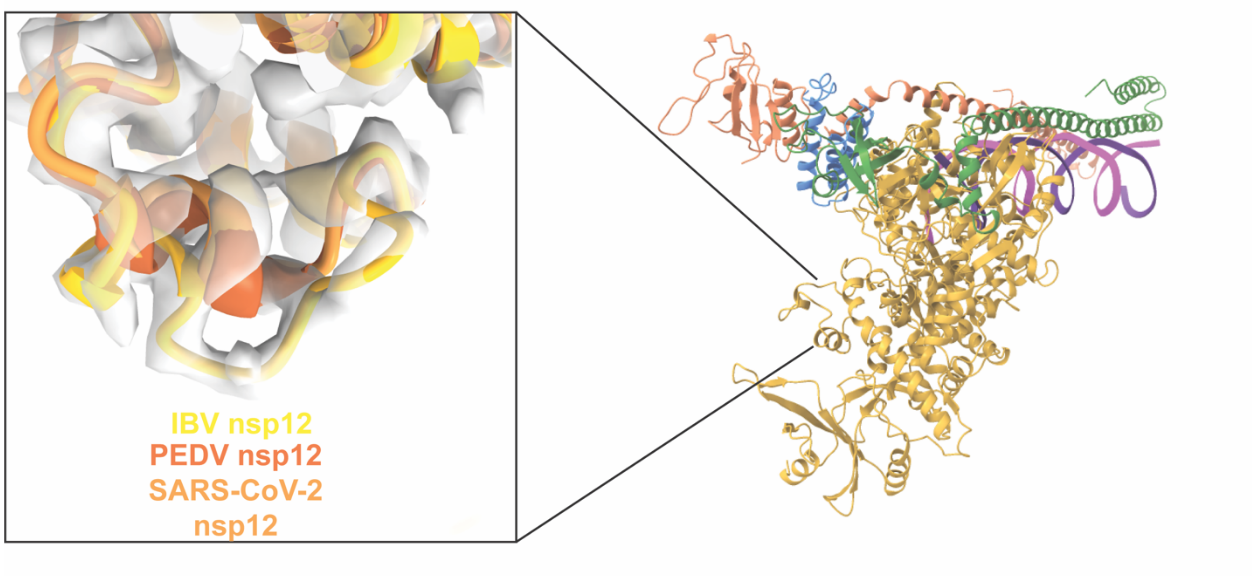


**Figure S14 Structural view of IBV nsp12 156-169 insertion:** Cartoon model of the IBV complex (right) with zoomed in view of superimposed models of IBV, PEDv (8URB) and SARS-CoV-2 (6XEZ*)*. Focused view on IBV nsp12 156-169 which has a 4 amino acid insertion. IBV model electron density shown to support altered loop conformation.

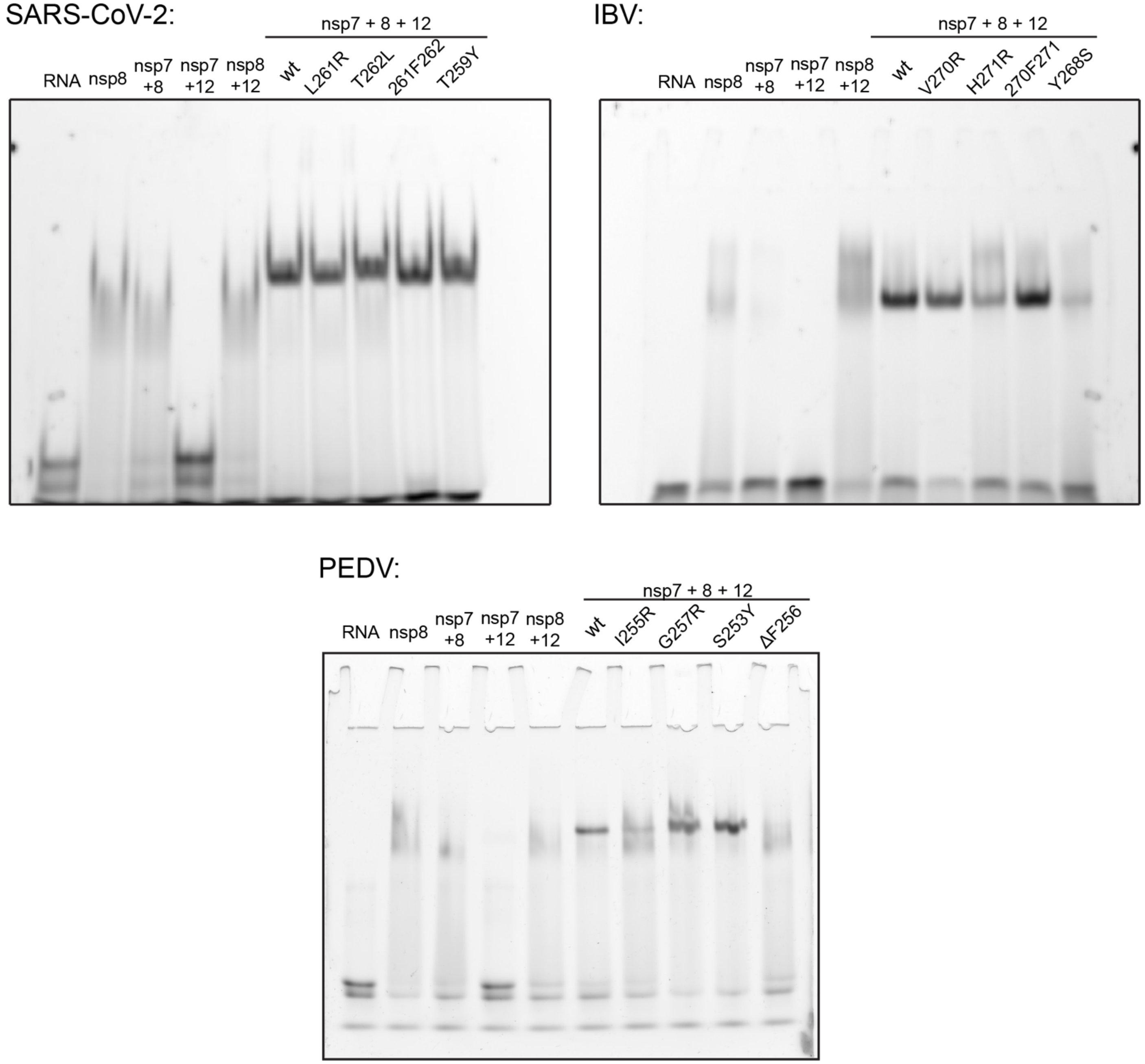


**Figure S15 Mutant nsp12 RTC RNA substrate affinity**: Native PAGE gel assessing RTC binding to FAM tagged RNA duplex with different mutant nsp12s for SARS-CoV-2, IBV, and PEDV.

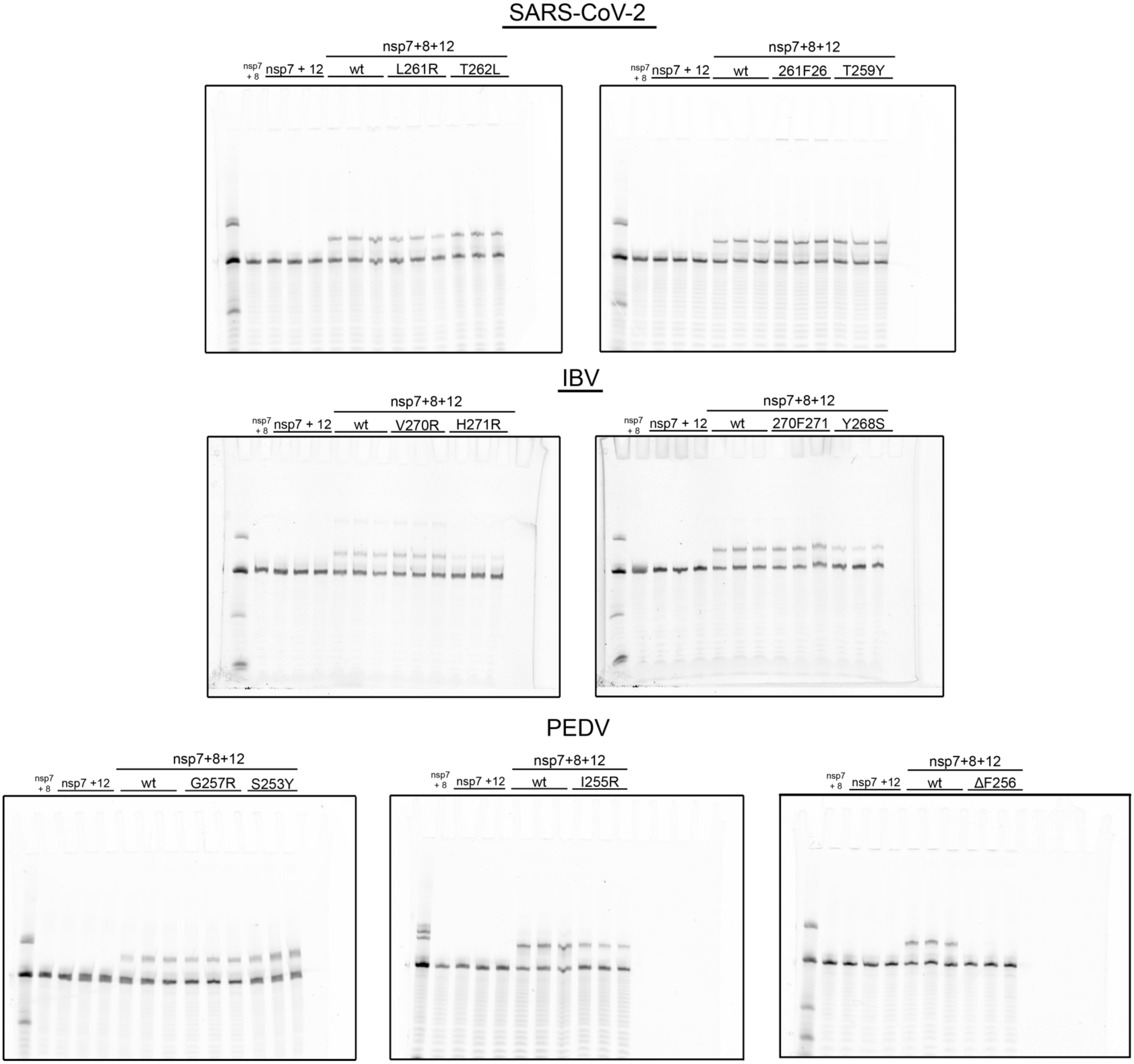


**Figure S16 Mutant nsp12 primer extension assays:** Primer extension assay assessing RTC activity with different mutant nsp12s for SARS-CoV-2, IBV and PEDv. Each gel contains the same piecewise controls of nsp7+8, nsp7+12 and wild type nsp7+8+12 performed in triplicate.
